# YAP-independent mechanotransduction drives breast cancer progression

**DOI:** 10.1101/495499

**Authors:** Joanna Y. Lee, Jessica Chang, Antonia A. Dominguez, Sungmin Nam, Julie Chang, Hong-pyo Lee, Sushama Varma, Lei S. Qi, Robert B. West, Ovijit Chaudhuri

## Abstract

Increased tissue stiffness is a driver of breast cancer progression. The transcriptional regulator YAP is considered a universal mechanotransducer, based largely on 2D culture studies. However, the role of YAP during *in vivo* breast cancer remains unclear. Here, we find that mechanotransduction occurs independently of YAP in breast cancer patient samples and mechanically tunable 3D cultures. Mechanistically, the lack of YAP activity in 3D culture and *in vivo* is associated with the absence of stress fibers and an order of magnitude decrease in nuclear cross-sectional area relative to 2D culture. This work highlights the context-dependent role of YAP in mechanotransduction, and establishes that YAP does not mediate mechanotransduction in breast cancer.

**One Sentence Summary:** Breast cancer mechanotransduction occurs independently of YAP, due to compact nuclear morphologies *in vivo*.

## Main Text

Enhanced mammographic density, associated with a ten-fold increase in extracellular matrix (ECM) stiffness, is one of the strongest risk factors for breast cancer progression (Fig. 1A) (1–6). Previous studies show that increased stiffness promotes a proliferative and invasive phenotype in mammary epithelial cells (7–10). During breast cancer progression, cancer cells “invade” through the basement membrane (BM) allowing metastatic dissemination to begin (11), resulting in decreased patient survival. Thus, there is a critical need to understand how enhanced ECM stiffness drives invasion.

**Fig. 1.**
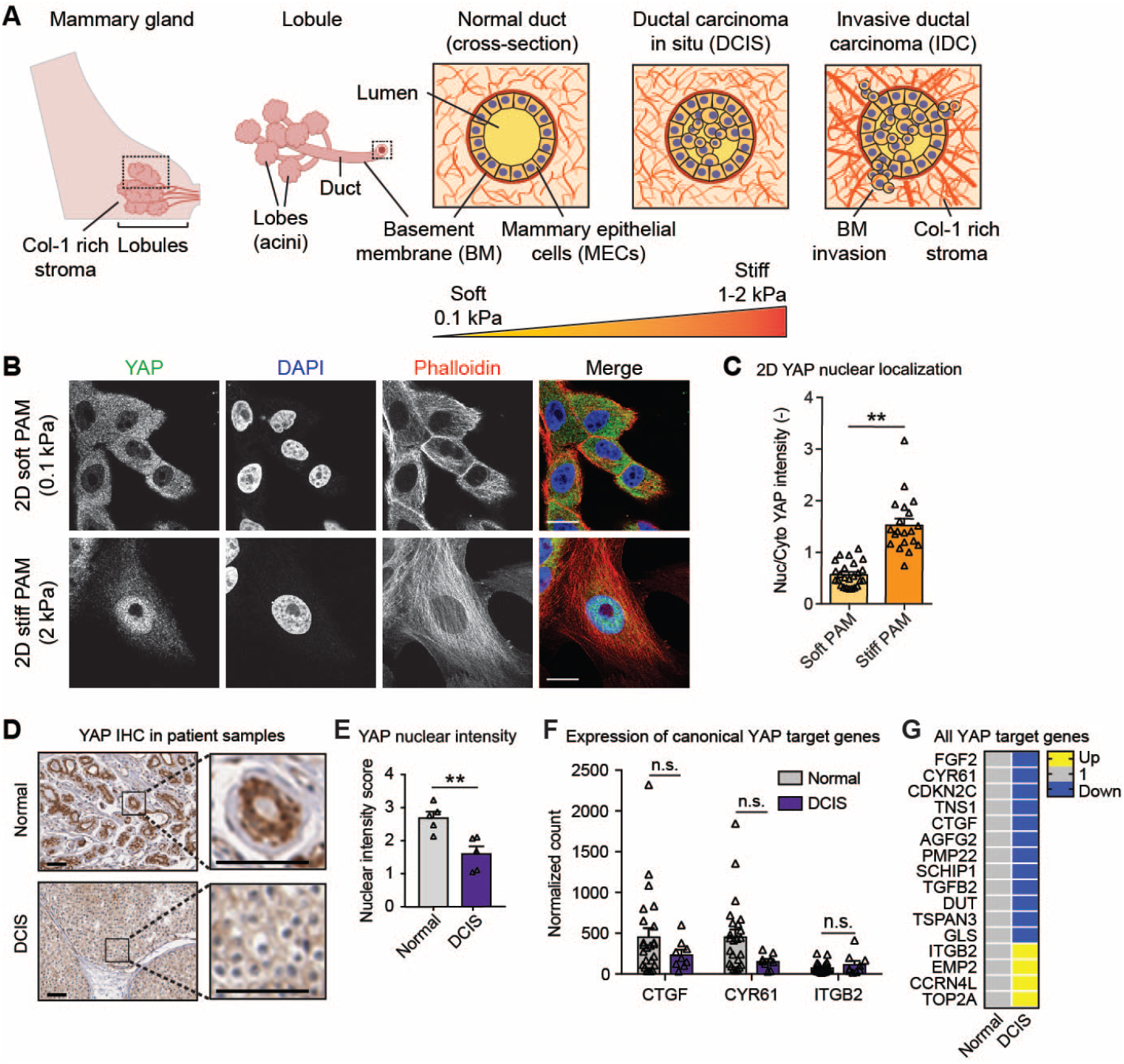
YAP is not activated during DCIS. **(A)** Schematic of DCIS progression. **(B)** MCF10A cells seeded on col-1-coated PAM gels. Bars: 10 μm. **(C)** YAP quantification from 2D gels. **(D)** YAP staining in primary tissues. Bars: 50 μm. **(E)** Quantification of YAP IHC intensity. Expression of **(F)** canonical and **(G)** all YAP target genes in patient samples.

YAP (Yes-associated protein), a transcriptional regulator that is deregulated in diverse cancers, has been implicated as a universal mechanotransducer (12). Mammary epithelial cells (MECs) cultured on increasingly stiff 2D polyacrylamide (PAM) gel substrates show YAP accumulation in the nucleus, activating expression of YAP target genes (Fig. 1B, C) (9,13). On stiff 2D substrates, stress fibers mediate flattening of the nucleus, which results in stretching of nuclear pores and YAP accumulation in the nucleus (14–16). However, cell morphology and signaling are significantly altered by culture dimensionality (17–20), and 3D culture has been reported to be crucially important when modeling breast cancer (19). In fact, several recent studies implicate YAP as a tumor suppressor during *in vivo* breast cancer (21–23). As such, the role of YAP in mechanotransduction during breast cancer is unclear.

To determine if YAP is responsible for mechanotransduction during breast cancer invasion we examined DCIS patient samples, a carcinoma state marked by increased ECM stiffness preceding BM invasion. Immunohistochemical (IHC) stains of patient samples show that YAP does not localize to the nucleus in DCIS samples (Fig. 1D, E). Additionally, 3SEQ analyses of patient samples for canonical YAP target genes (Fig. 1F) and additional YAP targets (Fig. 1G) show a lack of YAP activation. Further, analyses of publicly accessible gene expression datasets similarly showed no increase in YAP target gene expression with breast cancer (fig. S1A-E). However, expression of a subset of YAP target genes was increased in IDC samples (fig. S1F), which occurs post-BM invasion, suggesting that YAP activation may be relevant to post-invasion stages of breast cancer. Together, these analyses of three independent sets of patient data establish that YAP is not activated during early stages of breast cancer, when increased stiffness is reported to drive invasion.

We next examined whether YAP is responsive to increased ECM stiffness using a mechanically tunable 3D culture model of the mammary epithelium. Traditional mechanically tunable 3D culture models commonly incorporate col-1, which is highly relevant to post-invasion IDC (20). However, col-1 is not present in the BM and can activate tumorigenic signaling independently of stiffness (24, 25). Therefore, to mimic increased stiffness in a BM microenvironment without confounding col-1 signaling, we generated interpenetrating networks (IPNs) of reconstituted BM (rBM) with alginate (10). Addition of Ca^2+^ crosslinks the alginate network, increasing matrix stiffness without altering protein concentration, matrix architecture, or pore size (10). Elastic moduli of hydrogels ranged from ~0.04 kPa for “soft” to ~2 kPa for “stiff’ gels, covering the range of stiffness observed during breast cancer progression (Fig. 1A; see also fig. S2). We also generated traditionally used rBM and col-1 gels as controls (fig. S2). All hydrogels were used to encapsulate MCF10A cells, a non-transformed MEC line, in 3D culture acinar formation assays (fig. S2). Surprisingly, cells embedded in stiff IPNs or stiff col-1 gels, conditions that robustly promoted proliferation, invasion, and other markers of malignancy (i.e. β1 integrin and p-FAK) showed cytoplasmic YAP (Fig. 2A, B; see also fig. S3 and S4). Localization of YAP paralog TAZ mirrored YAP under all hydrogel conditions (fig. S5). Importantly, positive control experiments treating cells with nuclear export inhibitor Leptomycin B (LepB) showed strong YAP nuclear localization, similar to that of cells seeded on stiff 2D col-coated PAM gels, demonstrating that 3D cultured MCF10A cells are competent for YAP activation (Fig. 2C, D).

**Fig. 2.**
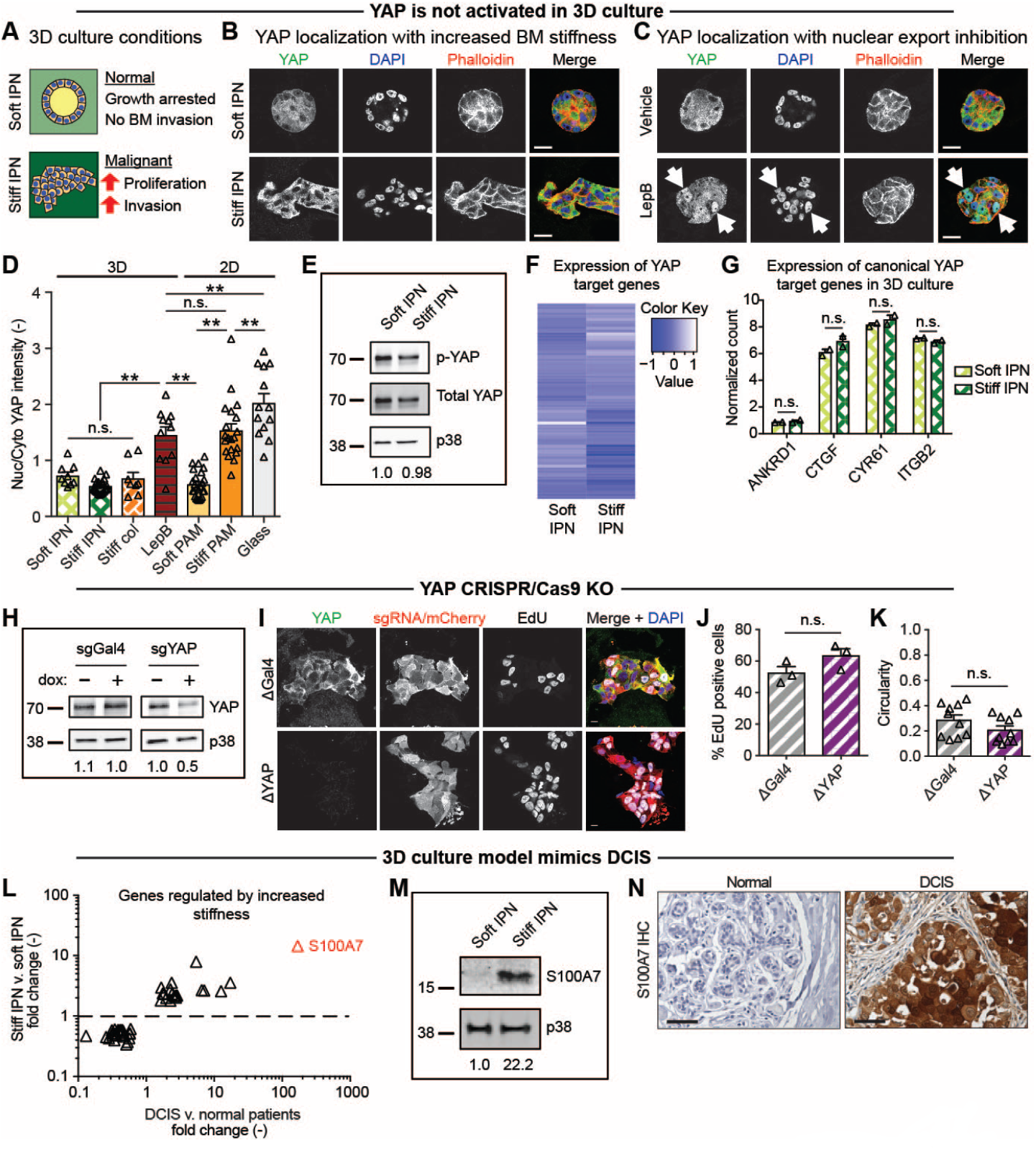
Enhanced ECM stiffness does not promote YAP activation in mammary epithelium in 3D culture. **(A)** Effects of 3D culture stiffness. **(B)** MCF10A cells encapsulated in 3D hydrogels. **(C)** Encapsulated cells treated withLeptomycin B (LepB). Arrows indicate nuclei with YAP. Bars: 20 μm. **(D)** YAP quantification from 3D and 2D (control) culture conditions. **(E)** Western blot analysis of p-YAP (S127) from 3D culture. p38 was used as a loading control. Quantification of bands (p-YAP/total YAP/p38) below each lane. **(F)** RNA-seq of YAP target genes (asidentified by Dupont et al., 2011) in 3D culture. **(G)** RNA-seq of canonical YAP target genes in 3D culture. **(H)** Western blot analysis of dox-inducible MCF10A::Cas9/sgGAL4 or sgYAP cells. Quantification of bands (YAP/p38) below each lane. **(I)** CRISPR/Cas9 cells encapsulated with dox. Bars: 10 μm. **(J)** Proliferation of cells from (I). **(K)** Invasiveness of cells from **(I)** as measured by cell cluster circularity. Only cells verified by IF for KO were assayed. **(L)** Set of genes regulated by enhanced stiffness in IPNs also upregulated in DCIS patient samples. Symbols represent each gene. Most highly enriched gene (S100A7) highlighted in red. **(M)** Western blot analysis of S100A7 from 3D culture. Quantification of bands (S100A7/p38) indicated below each lane. **(N)** S100A7 staining in primary tissue. Bars: 50 μm.

We explored the possibility that YAP activation requires a higher range of stiffness. Although 1-2 kPa stiff hydrogels are sufficient to induce proliferation and invasion (fig. S3) and are physiologically relevant for malignant mammary tissue (4, 5), we generated 20 kPa hydrogels, an order of magnitude stiffer than malignant mammary tissue. 20 kPa gels similarly failed to induce nuclear localization of YAP in MCF10A cells (fig. S6A, B). Additionally, increasingly malignant MEC lines, MCF10AT and MCF10CA1a (26), and preformed acinar structures transplanted into stiff hydrogels also did not display increased YAP nuclear localization (fig. S6C-D).

In addition to examining YAP localization, we assessed YAP activity through analysis of YAP phosphorylation and gene expression. Western blot (WB) analysis of cells harvested from soft and stiff IPNs showed similar levels of YAP S127 phosphorylation, a mark of cytoplasmic retention and thus inactivity (Fig. 2E). RNA-seq was next performed to assay expression of YAP transcriptional targets, using the YAP target gene list used to identify YAP as a mechanotransducer in 2D culture (table S1) (13). In agreement with IF results, expression of YAP target genes did not trend with increased stiffness (Fig. 2F; see also table S1) or col-1 density (fig. S7A). Notably, expression of canonical YAP target genes ANKRD1, CTGF, CYR61, ITGB2 were not differentially regulated by enhanced stiffness (Fig. 2G) or col-1 density (fig. S7B). This is in contrast to the robust YAP activation and target gene expression demonstrated by the same MEC line in 2D culture (fig. S8) (27).

Given the surprising lack of YAP activation by increased 3D culture stiffness, we generated doxycycline (dox)-inducible CRISPR/Cas9 YAP knockout (ΔYAP) MCF10A cells to test the dispensability of YAP in mechanotransduction. As absence of YAP may impact cell growth in 2D culture prior to encapsulation, cells stably expressing dox-inducible Cas9 and sgRNA targeting YAP (MCF10A::Cas9/sgYAP) were first encapsulated in 3D culture without Cas9 induction. Following encapsulation, cells were treated with dox to induce Cas9 expression and sgRNA-directed editing. Dox treated MCF10A::Cas9/sgYAP cells showed depletion of YAP protein compared to untreated and MCF10A::Cas9/sgGAL4 controls (Fig. 2H). As Cas9 induction results in a mixed population of KO cells, only cells verified for ΔYAP by IF were assayed for mechanotransduction (Fig. 3I). Interestingly, ΔYAP cells did not reduce stiffness-induced invasion (Fig. 2J) or proliferation (Fig. 2K) compared to ΔGAL4 controls. As YAP did not regulate mechanotransduction during breast cancer progression, we explored other transcriptional regulators that target genes identified by RNA-seq to be modulated by stiffness (fig. S9-11). Bioinformatics, small molecule inhibitor, inducible CRISPR/Cas9 KO, and overexpression experiments strongly implicate STAT3 and p300 as mechanotransducers during breast cancer (fig. S9 and S10). Taken together, our analyses of YAP and TAZ nuclear localization, YAP phosphorylation state, expression of YAP target genes, and inducible CRISPR/Cas9 knockout cells conclusively show that YAP does not mediate mechanotransduction in 3D culture.

**Fig. 3.**
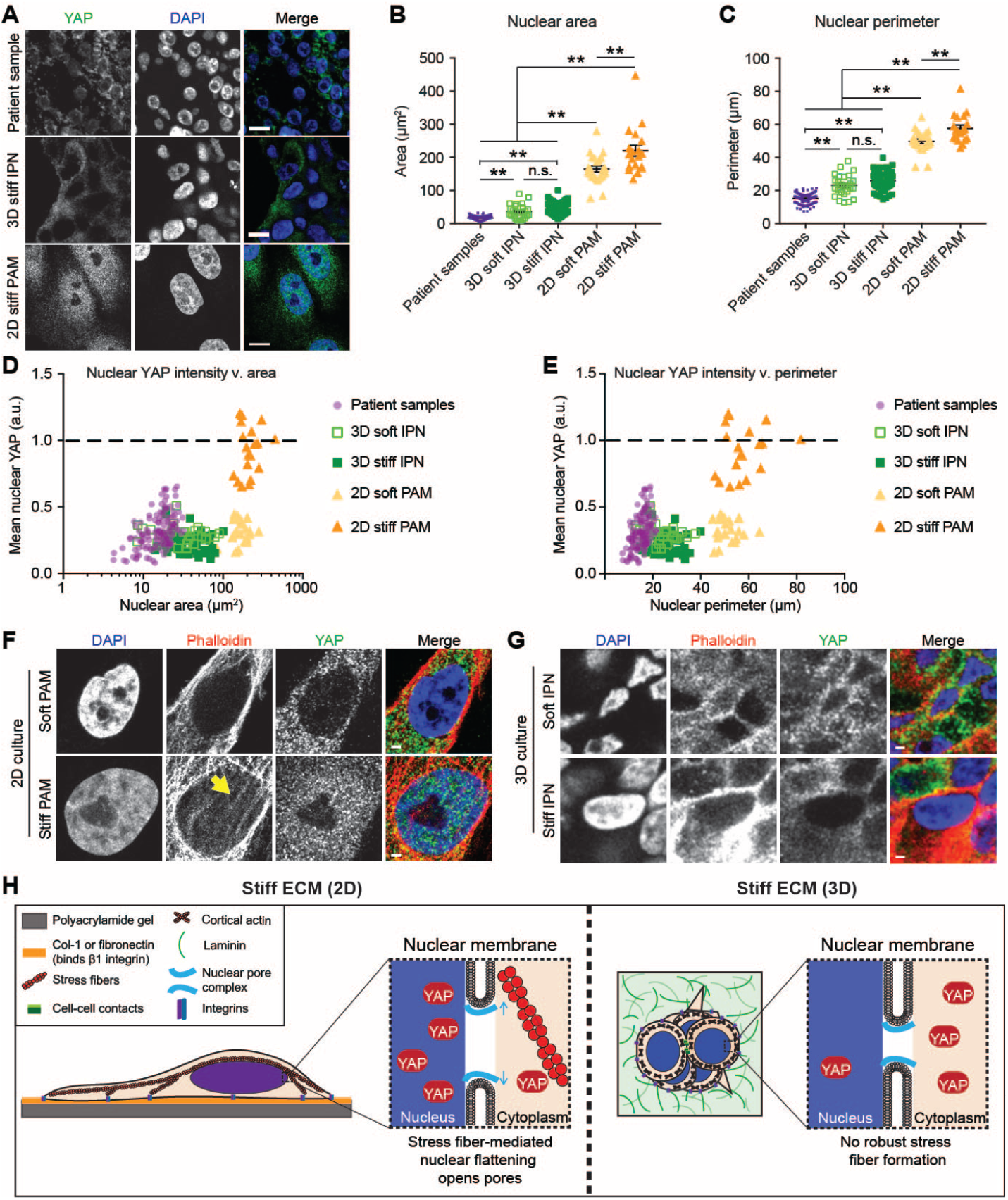
Nuclear morphologies, which drive YAP localization, are distinct in 2D culture from 3D culture and *in vivo*. **(A)** Images of nuclear morphologies and YAP. Bars: 10 μm. **(B)** Areas and **(C)** perimeters of nuclei. Patient samples from five DCIS patients. YAP intensity with nuclear **(D)** area and **(E)** perimeter. Values normalized by positive controls within each sample. Dotted line represents positive nuclear YAP. F-actin staining in **(F)** 2D or **(G)** 3D culture. Arrow indicates perinuclear stress fibers. Bars: 2 μm. **(K)** Model of stiffness-induced YAP localization in 2D v. 3D.

To assess the relevance of this 3D culture model to DCIS, we compared our RNA-seq data of cells encapsulated in soft or stiff IPNs (Fig. 2F) to 3SEQ data from normal and DCIS patient samples (28) (Fig. 2F, G). Importantly, a set of genes was identified that showed similar regulation in stiff IPNs as DCIS samples (Fig. 2L; see also and fig. S11 and table S2). Interestingly, RNA-seq of cells isolated from stiff col-1 containing gels show a distinct gene expression profile compared to BM stiffness, and captures key aspects of the gene expression profile in IDC patient samples (fig. S11). Plotting fold change *in vitro* (i.e. stiff IPNs) against fold change *in vivo* (i.e. DCIS patient samples) revealed the most highly upregulated target from stiff IPNs, S100A7, as the most relevant stiffness-regulated gene in DCIS (Fig. 5L). S100A7 has been implicated in DCIS with roles in proliferation and apoptosis-resistance, and tumor-associated immune cell recruitment (29–31). RNA-seq results were confirmed by WB analysis of S100A7 in cells harvested from soft and stiff IPNs (Fig. 2M), IHC of S100A7 in breast cancer patient tissues (Fig. 2N), and qPCR of cells harvested from soft and stiff IPNs (fig. S12). Together, these results demonstrate that 3D culture of MECs in stiff IPNs is highly relevant to modeling DCIS, and provides a gene signature of stiffness-induced carcinoma progression.

To elucidate the mechanism underlying the confounding result that YAP is responsible for mechanotransduction in 2D, but not 3D culture nor primary tissue, we examined nuclear morphologies. This analysis was motivated by the recent finding that stiffness-induced YAP activation requires nuclear flattening-mediated opening of nuclear pores (15, 16). Analysis of nuclear morphologies showed drastic differences in DCIS primary tissues and cells in 3D culture compared to 2D culture (Fig. 3A). Strikingly, nuclear area in cells from 2D culture show a ten-fold increase in cross-sectional area compared to 3D culture and patient samples (Fig. 3B). A three-fold increase in nuclear perimeter was also observed (Fig. 3C), in addition to a significant increase in solidity, a measure of the “smooth” nature of the perimeter (fig. S13A). During progression from normal to DCIS to IDC, patient samples showed interesting, but comparatively small, differences in nuclear morphology (Fig. S13B-E).

Notably, YAP nuclear intensity scales with nuclear area, with nuclei from patient samples and 3D culture deviating from the size range observed for positive nuclear YAP intensity (Fig. 3D). Similarly, nuclear YAP intensity scales with nuclear perimeter (Fig. 3E). Positive nuclear YAP intensity also correlated with high solidity, which is almost exclusively observed in nuclei from 2D culture (Fig. S13F). However, some nuclei from soft 2D PAM reach the required size ranges but fail to show positive nuclear YAP intensity, suggesting that nuclear morphology is not the only factor required for YAP activation.

As nuclear morphologies and YAP activation in 2D culture has been linked to stress fiber contractility, we investigated the role of stress fibers in 3D culture. Recent studies showed mechanical coupling of stiff ECM to the nucleus through stress fibers, with fiber contractility causing nuclear flattening and subsequent nuclear pore stretching (15, 16). Further, the enrichment of perinuclear stress fibers was required for YAP nuclear translocation in 2D culture (16). To examine if stress fibers contribute to the observed changes in nuclear morphology, we assayed stress fibers in cells cultured in 2D and 3D. The presence of robust perinuclear stress fibers was observed in cells cultured on stiff PAM gels, in which nuclear localization of YAP was observed, but not soft PAM gels nor stiff or soft IPNs, in which nuclear localization of YAP was not observed (Fig. 3F, G). This is in agreement with previous reports that cells cultured in 3D substrates fail to form robust stress fibers (32, 33), and instead adopt a predominantly cortical F-actin architecture (Fig. 3G). This suggests a model where the presence of perinuclear stress fibers, coupled with distinct nuclear morphologies, is the basis of differences between YAP 2D and 3D activation (Fig. 3H).

In this study, we examined the role of YAP in mediating mechanotransduction during ductal carcinoma progression using patient samples and 3D culture models. Cancer has historically been thought of as a genetic disease, with tumors arising from genetic mutations in DNA. However, it has been increasingly recognized that the microenvironment plays a key role in regulating cancer progression. Our study provides compelling evidence that 2D YAP mechanotransduction studies do not recapitulate the conditions seen in clinical samples, and suggests a critical need for the use of 3D culture models in studying breast cancer. Finally, our findings also reveal new therapeutics targets, including STAT3 and p300, for blocking invasion in breast cancer.

## Acknowledgments

We gratefully acknowledge Katrina Wisdom (Stanford University) and Ryan Stowers (Stanford University) for assistance with mechanical testing, materials, and helpful discussions. We acknowledge Robert Blake (Genentech) for helpful advice and assistance with p300 inhibitors. We thank Lalage Wakefield and Binwu Tang (NIH) for assistance with MCF10AT and MCF10CA1a cell lines. We thank Bayana Science and Sasha Denisin (Stanford University) for careful reading of the manuscript.

## Author contributions

Conceived and designed the experiments: JYL OC. Performed the experiments: JYL JC AD JC HL SV. Analyzed the data: JYL JC OC. Contributed reagents/materials/analysis tools: JYL JC AD SN RW OC. Supervised and administered the project: LSQ RW OC. Acquired funding: OC. Wrote the paper: JYL OC.

## Data and materials availability

RNA-seq data are stored in GEO with the accession code GSE102506. 3SEQ breast cancer progression data are available as described in the original manuscript (28).

## Supplementary Materials

### Materials and Methods

Step-by-step protocols can be accessed at The Chaudhuri Lab’s bio-protocol website: https://en.bio-protocol.org/bio101/labinfo.aspx?labid=40&lang=0

#### Cell culture and cell lines

MCF10A cells obtained from the ATCC (cat. #CRL-10317; ATCC) were cultured in DMEM/F12 50/50 medium (cat. #11330057; Thermo Fisher Scientific) supplemented with 5% horse serum (cat. #16050122; Thermo Fisher Scientific), 20 ng/ml EGF (cat. #AF-100-15; Peprotech, Inc.), 0.5 μg/ml hydrocortisone (cat. #H0888-1G; Sigma), 100 ng/ml cholera toxin (cat. #C8052-1MG; Sigma), 10 μg/ml insulin (cat. #91077C-250MG; Sigma), and 100 U/ml Pen/Strep (cat. #15140; Thermo Fisher Scientific) as previously described (1). MCF10AT and MCF10CA1a cells were a gift from Lalage Wakefield (NIH) and were cultured in complete medium for experimental consistency.

For inducible MCF10A::Cas9 cell line, lentivirus was produced harboring Edit-R Inducible Lentiviral hEF1α-Blast-Cas9 Nuclease Plasmid DNA (cat. #CAS11229; Dharmacon) (see Cloning and lentiviral generation below). Following infection, Cas9 cells were maintained in MCF10A growth medium as above, supplemented with 5 μg/ml blasticidin (cat. #R21001; Thermo Fisher Scientific). Following a second round of infection with indicated sgRNAs (see Cloning and lentiviral generation below)., MCF10A::Cas9/sgRNA cell lines were maintained in medium supplemented with 5 μg/ml blasticidin and 1 μg/ml puromycin (cat. #A1113803; Thermo Fisher Scientific).

#### Reagents

EdU incorporation assay (cat. #C10337; Thermo Fisher Scientific) was performed according to manufacturer’s instructions with a 24 h incubation of 10 μM EdU.

For inhibitor studies, MCF10A cells encapsulated in hydrogels were incubated in Genentech p300 inhibitor GNE-049 at a final concentration of 0.5 μM (Genentech, Inc.; MTA OR# 216339) or STAT3 peptide inhibitor PY*LKTK at a final concentration of 500 μM (cat. #ab142104; Abcam). p300 inhibitor C646 (cat. #SML0002; Sigma) was also used at the indicated concentrations.

CRISPR/Cas9 MCF10A cell lines were generated by first producing a doxycycline-inducible Cas9/blast MCF10A cell line. Cas9/blast transfer vector (cat. #CAS11229; Dharmacon) containing virus was produced and used to infect WT MCF10A cells. Stably expressing cells were selected using 5 μg/ml blast. MCF10A::Cas9/blast cells were infected with lentivirus harboring sgRNA against GAL4/mCherry/puro, YAP/mCherry/puro, or STAT3/mCherry/puro. Doubly stably-expressing cells were selected using 5 μg/ml blast with 1 μg/ml puro. CRISPR/Cas9 editing was induced by adding 2 μg/ml dox (cat. #AAJ6042206; Alfa Aesar) and knockout verified by WB and IF.

#### Cloning and lentiviral generation

| Addgene ID: | Plasmid name: |
| --- | --- |
| 121423 | pLenti-sgYAP-2 |
| 121424 | pLenti-sgYAP-10 |
| 121425 | pLenti-sgSTAT3-1 |
| 121426 | pLenti-EGFP |
| 121427 | pLenti-STAT3-linker-EGFP |

sgRNAs were expressed using a lentiviral mouse U6 (mU6) promoter-driven expression vector that coexpessed Puro-T2A-mCherry from a CMV promoter. sgRNA sequences were generated by PCR and introduced by InFusion cloning into the sgRNA expression vector digested with BstXI and XhoI. sgSTAT3-1 sequence: TCAGGATAGAGATAGACCAG. For YAP, 2 sgRNA sequences were used and pooled during lentiviral production. sgYAP-2 sequence: AGATGACTTCCTGAACAGTG; sgYAP-10 sequence: GAATCACCCTGAGTCAGGAG.

To assemble pLenti-STAT3-linker-EGFP for overexpression, STAT3 was amplified from pLEGFP-WT-STAT3, with the forward primer containing the linker sequence, and inserted using Infusion Cloning into MluI and EcoRI digested pLenti-Origene-Nrf21. To assemble pLenti-EGFP control, EGFP was amplified from pLenti-Origene-Nrf21 and inserted into XhoI and EcoRI digested pLenti-Origene-Nrf21.

For lentiviral generation, HEK293T cells were seeded at 1 x 107 cells/10 cm dish. The next day 70-90% confluent cells were transfected. For each dish, 9 ug of lentiviral transfer vector, 8 μg of packaging vector pCMV-dR8.91 and 1 ug of packaging vector pMD2-G were transfected using Lipofectamine 3000 Transfection Reagent (cat. #L3000008; ThermoFisherScientific) Opti-MEM Reduced Serum Medium (cat. #31985062; Gibco) according to the manufacturer’s instructions. Medium was replaced with complete medium 4 h following transfection. 48 h following transfection, lentivirus-containing supernatant was harvested and filtered through a 0.22 μm Steriflip (Millipore). Filtered supernatant was concentrated using Lentivirus Precipitation Solution (cat. #VC100; AlStem) according to the manufacturer’s instructions. Following concentration, lentiviral pellets were resuspended in 1/100 of original volume using cold DMEM/F12 and stored at −80°C. For MCF10A transduction, concentrated lentivirus was added to complete medium containing 8 μg/ml polybrene (cat. #sc134220; Santa Cruz Biotech) at a volume of 1:100.

#### Hydrogel formation

Matrigel (cat. #354230; Corning) was purchased for use as rBM matrix and used at a final concentration of 4.4 mg/ml for all experiments. Collagen-1, derived from rat tail, (cat. #354236; Corning) was lyophilized and reconstituted in 20 mM acetic acid. Immediately before cell encapsulation, reconstituted col was supplemented with 10x PBS, neutralized with 0.1 M NaOH, and pH adjusted with 0.1 N HCl. rBM and col were mixed with cells and DMEM/F12 to the reach the indicated final concentrations. MCF10A cells were trypsinized, strained through a 40 μm cell strainer to enrich for single cells, counted on a Vi-CELL (Beckman Coulter Life Sciences), and seeded at a final concentration of 1 x 105 cells/ml hydrogel. Hydrogel-cell mixtures were quickly deposited into wells of a 24-well plate pre-coated with 50 μl gelled rBM. Hydrogels containing cells were placed in a 37°C incubator with CO2 to gel for 30 min before a transwell insert (Millipore) was placed on top to prevent floating and 1.5 ml complete medium added.

IPNs were formed as described (2). Briefly, LF20/40 alginate (FMC Biopolymer) was solubilized, dialyzed, charcoal filtered, sterile filtered, lyophilized, and reconstituted to 2.5% w/v in DMEM/F12. Alginate was mixed with rBM, cells, and DMEM/F12 and loaded into a 1 ml Luer lock syringe (Cole-Parmer), on ice. For crosslinking, a second 1 ml syringe was loaded with 125 mM CaSO_4_ or DMEM/F12, on ice. Syringes were connected with a female-female Luer lock coupler (ValuePlastics), rapidly mixed with 4-6 pumps of the syringes handles back and forth, and quickly deposited into pre-coated wells, as above. IPNs containing cells were allowed to gel before adding transwell filters and medium, as above.

For 20 kPa alginate-RGD hydrogels, alginate-RGD hydrogels were prepared as described previously (3). Briefly, LF20/40 alginate (FMC Biopolymer) was dialyzed, filtered and lyophilized, and then was coupled to RGD oligopeptide GGGGRGDSP (Peptide 2.0) using carbodiimide chemistry (4). The final density of RGD in the alginate hydrogel was matched as 150 mM RGD in a 2% wt/vol alginate gel. The modified alginate was dialyzed, charcoal filtered, sterile filtered and lyophilized again. Alginate-RGD was reconstituted to 2.5% w/v in DMEM/F12 and mixed with MCF10A cells. The cell-alginate solution was then mixed with DMEM/F12 containing 24.4 mM CaSO_4_, and then deposited between two glass plates spaced 2 mm apart. The cell-alginate mixture was allowed to gel for 45 minutes, and then disks of hydrogel were punched out and immersed in complete growth medium.

For 2D PAM gels, the surface of coverslips was functionalized accoring to a previous method (5). Coverslips were cleaned with ethanol, immersed in 0.5% (3-Aminopropyl)trimethoxysilane (in dH_2_O) at room temperature for 30 min and washed with dH_2_O. Coverslips were then immersed in 0.5% glutaraldehyde in dH_2_O at room temperature for 30 min and dried. A prepolymer solution was prepared containing acrylamide, N,N’-methylene-bis-acrylamide, 1/100 volume of 10% Ammonium Persulfate (APS), and 1/1000 volume of N,N,N’,N’-Tetramethylethylenediamine (TEMED). The final concentration of acrylamide and bis-acrylamide was varied to control substrate stiffness (6). For 0.1 kPa hydrogels, 3% /0.02% were used. For 1 kPa hydrogels, 3% /0.1% were used. For 2 kPa hydrogels, 4% /0.1% were used. Prepolymer solutions were deposited on a Sigmacote-treated hydrophobic glass plate, and functionalized coverslips placed on top of the prepolymer solution. Polyacrylamide solutions were allowed to polymerize for 30 minutes between the hydrophobic glass plate and the functionalized coverslip. When polymerization was completed, polyacrylamide gels were carefully separated from the glass plate.

To enable cell adhesion to the PAM gel, col-1 and rBM were conjugated to the gel surface using sulfosuccinimidyl 6-(4’-azido-2’-nitrophenylamino)hexanoate (sulfo-SANPAH) as a protein-substrate linker. PAM gels were incubated in 1 mg/ml sulfo-SANPAH in 50 mM HEPES pH 8.5, activated with UV light (wavelength 365 nm, intensity 4mW/cm2) for 20 min, washed in HEPES, and then incubated in 100 ug/ml col-1 and rBM in HEPES overnight at room temperature. The protein-crosslinked PAM gels were washed with PBS before use.

#### Mechanical Testing

Stiffness measurements of 3D culture rBM, col, and IPN hydrogels were performed using an AR-G2 stress-controlled rheometer with 25-mm top- and bottom-plate stainless steel geometries (TA Instruments). Hydrogel solutions without cells were mixed and immediately deposited onto the bottom plate of the rheometer and the top plate lowered such that the gel formed a uniform disk between the two plates. Exposed hydrogel surfaces were coated with mineral oil (Sigma) and covered with a hydration chamber to prevent sample dehydration. The storage modulus was monitored at 37°C with 1% strain at a frequency of 1 Hz and measurements taken once the storage modulus reached an equilibrium value. The storage and loss moduli were then used to calculate the Young’s modulus (E). Young’s moduli (i.e. elastic moduli) were calculated using the equation *E* = 2*G\** × (1 + *v*), where *v* is Poisson’s ratio, assumed to be 0.5, and *G\** is the bulk modulus calculated using the equation *G\** = (*G’^2^* + *G’’^2^*)^1/2^ where *G’^2^* is the storage and *G’’^2^* is the loss modulus.

To measure substrate stiffness of 2D PAM gels, unconfined compression tests were performed using an Instron MicroTester 5848. PAM gels were compressed at a rate of 1 mm/min. The tangent elastic modulus of the measured stress-strain curves was calculated between 5%-15% strain (7, 8). Stiffness of 3D culture alginate hydrogels was measured using unconfined compression tests according to a previously published method (3). Briefly, alginate disks (15 mm in diameter, 2 mm thick) were submerged in DMEM for 1 day to fully equilibrate. The gel disks were compressed to 15% at a rate of 1 mm/min and the slope of the stress-strain curve from 5% to 10% strain was used to obtain the stiffness of alginate hydrogel.

#### Antibodies

Mouse anti-YAP (cat. #sc-101199; Santa Cruz Biotech) was used at 1:200 (IF) and 1:500 (WB), rabbit anti-phospho-YAP (cat. #13008; Cell Signaling Technology) was used at 1:500 for WB. Mouse anti-S100A7 (cat. #sc-377084; Santa Cruz Biotech) was used at 1:500 (WB). Mouse anti-S100A7 (cat. #HPA006997; Millipore-Sigma) was used at 1:100 for IHC. Rabbit anti-phospho p300 (cat. #ab135554; Abcam) was used at 1:500 for WB, mouse anti-p300 (cat. #sc-32244; Santa Cruz Biotech) was used at 1:500 for WB. Rabbit anti-phospho STAT3 (cat. #ab76315; Abcam) was used at 1:500 for WB, mouse anti-STAT3 (cat. #sc-8019; Santa Cruz Biotech) was used at 1:500 for WB. Rabbit anti-p38 (cat. #sc-535; Santa Cruz Biotechnology) was used at 1:2000 for WB. Mouse anti-β1 integrin (cat. #ab24693, Abcam) was used at 1:500 for IHC. Rabbit anti-phospho FAK (cat. #31H5L17; Thermo Fisher Scientific) was used at 1:100 for IHC. Rabbit anti-α-actinin (cat. #3134; Cell Signaling Technology) was used at 1:500 for WB. Rabbit anti-β-actin (cat. #8457; Cell Signaling Technology) was used at 1:1000 for WB. Phalloidin-Alexa555 (A34055; Thermo Fisher Scientific) was used at 1:100 and DAPI (cat. #D9542; Sigma-Aldrich) was used at 5 μg/ml for IF.

For IF, Alexa 488-, 555- or 647-conjugated secondary antibodies (Thermo Fisher Scientific) were used at 1:500. For WB, IRDye 680 or 800-conjugated secondary antibodies (LI-COR Biotechnology) were used at 1:10,000.

#### Western blotting and immunoprecipitation

MCF10A cells encapsulated for 7 days were harvested from IPNs by incubation in cold PBS containing 50 mM EDTA (Sigma) for 5 min while pipetting to break up gels. Cells were centrifuged at 500 x g for 10 min. The supernatant was removed and the cells with remaining matrix material were treated with 0.25% trypsin (Gibco) for 5 min and centrifuged for 5 min at 500 x g. Cell pellets were washed with 20% serum-containing resuspension buffer to neutralize trypsin and washed twice with PBS. For SDS-PAGE of whole cell lysates, MCF10A cells were lysed in Pierce RIPA buffer (cat. #89900; Thermo Fisher Scientific) supplemented with Protease Inhibitor Cocktail Tablets (cat. #11836170001; Roche) and PhosSTOP Phosphatase Inhibitor Cocktail Tablets (cat. #04906845001; Roche) according to the manufacturer’s instructions. Protein concentration was determined using the Pierce BCA Protein Assay Kit (cat. #23227; Thermo Fisher Scientific). Laemmli sample buffer (cat. #1610747; Bio-Rad) was added to lysates and samples boiled for 10 min before loading 25 μg protein in each lane of a 4-15%, 15- well, gradient gel (cat. # 4561086; Bio-Rad). Proteins were transferred to nitrocellulose at 100 V for 105 min, blocked with 5% milk in TBS-T (137 mM NaCl, 2.7 mM KCl, 19 mM Tris base, 0.1% Tween, pH 7.4), incubated in primary antibody overnight, IRDye 680- or 800-conjugated secondary antibodies (Li-COR Biotechnology) for 1 h, and visualized with the Li-COR Odyssey imaging system (Li-COR Biotechnology). Quantitative analysis of western blots was performed using the Li-COR Odyssey software (LI-COR Biotechnology).

For IPs, MCF10A cells were harvested from IPNs as above and lysed in Pierce IP buffer (cat. #87787; Thermo Fisher Scientific) containing inhibitors as above and quantified as above. 1 μg of antibodies were conjugated to 10 μl Dynabeads (cat. # 10001D; Thermo Fisher Scientific), added to cell lysate, and rotated for 30-60 min on ice. Beads were washed 3x with IP buffer containing inhibitors followed by elution of protein complexes off beads with sample buffer.

#### Immunofluorescence

Cells encapsulated in hydrogels for seven days were fixed for 30 min in 4% paraformaldehyde in DMEM/F12. Gels containing cells were washed with PBS and incubated in 30% sucrose in PBS with calcium and magnesium overnight followed by incubation in 50/50 30% sucrose/OCT for 6 h. Gels were embedded in OCT and frozen prior to cutting 40 μm sections using a Microm HM 550 Cryostat. Sections were blocked in PBS-BT+: PBS pH 7.4 (Gibco) supplemented with 1% BSA (Sigma-Aldrich), 0.1% Triton X-100 (Sigma-Aldrich), 0.3 M glycine (Sigma-Aldrich), 10% goat serum (Gibco), and 0.05% sodium azide (Sigma-Aldrich). Sections were incubated in primary antibodies diluted in blocking solution as indicated in “antibodies” section for 1 h, and then Alexa 488-, 555- or 647-conjugated secondary antibodies (Thermo Fisher Scientific) diluted 1:500 in blocking solution for 30 min. Sections of gels containing cells were imaged using a Leica TCS SP8 confocal laser scanning microscope (Leica Microsystems, Inc.) with an HC PL APO 63x (1.40 NA Oil CS2) objective. Images were collected from HyD and PMT detectors using LasX software and processed using Photoshop (Adobe Systems).

For morphology analyses, ImageJ was used to trace cell clusters and circularity measured using the Measurements function. Circularity, C, was calculated as, *C =* 4π(A/p2), where *A* is the area and *P* is the perimeter. A perfect circle would have a circularity of 1. Solidity was calculated as area enclosed by outer contour of object divided by area enclosed by convex hull of outer contour.

Cell Profiler was used to quantify YAP nuclear/cytoplasmic intensity in IF images. ImageJ was used to trace cell nuclei using DAPI images using the following macro (pixel/μm of image was first determined and replaced in “Set Scale” distance; found pixel/μm of image by drawing line over scale bar embedded in image and using the function Analyze -> Set Scale). Doublets or cell debris were then manually excluded. Nuclear traces were then overlaid on YAP images to measure mean nuclear YAP intensity using the following macro.

~~~
Macro to trace cell nuclei:
run(“Set Scale…”, “distance=[3.45] known=1 pixel=1 unit=μm”);
run(“Gaussian Blur…”, “sigma=2”);
run(“Subtract Background…”, “rolling=50”);
setAutoThreshold();
//run(“Threshold…”);
setAutoThreshold();
setThreshold(55, 255);
run(“Convert to Mask”);
run(“Fill Holes”);
run(“Watershed”);
run(“Find Edges”);
run(“Analyze Particles…”, “size=100-Infinity pixel circularity=0.00-1.00 show=Nothing exclude
add”);
close();

Macro to measure nuclear YAP intensity:
run(“Set Measurements…”, “area mean center perimeter bounding shape integrated skewness
redirect=None decimal=3”);
run(“Set Scale…”, “distance=[3.45] known=1 pixel=1 unit=μm”);
setOption(“Show All”,true);
roiManager(“Measure”);
saveAs(“Measurements”, “/Users/Joanna/Desktop/Results.xls”);
~~~

#### Tissue immunohistochemistry

Immunohistochemical (IHC) staining was performed on paraffin-embedded tissue microarray (TMA) sections (Stanford TA419, 445, 424). Anti-S100A7 polyclonal antibody at 1:100 (Millipore-Sigma St. Louis, MO, catalog # HPA006997) was used as primary antibody for IHC staining. Antibody was diluted in PBS. Briefly, 4 μm TMA sections were deparaffinized in 3 changes of xylene for 10 mins each and hydrated in gradient series of ethyl alcohol. Following target retrieval in 10 mM citrate pH6 (Dako/Agilent, Carpinteria, CA, USA, catalog #S2369) to retrieve antigenic sites at 116°C for 3 minutes. Staining was then performed using the VectaStain ABC anti-rabbit kit (Vector Laboratories Burlingame CA, USA, catalog #PK6101). Diaminobenzidine (DAB) (DAKO/Agilent, Carpinteria, CA, USA, catalog #K3468) was used at room temperature for 10 min for color development. The IHC Profiler macro for ImageJ was used to quantify YAP and S100A7 intensity in IHC samples 78.

#### RNA extraction and next generation sequencing

Gels containing MCF10A cells were frozen in liquid nitrogen, ground, and treated with ice-cold PBS supplemented with 50 mM EDTA to break up IPNs. RNA was harvested using a combination of TRIZOL reagent and GenCatch Total RNA Extraction Kit (Epoch). RNA-seq experiments were performed in biological replicate and cDNA libraries constructed using the TruSeq RNA Library Prep Kit v2. Libraries were sequenced on a single lane of the Illumina Hiseq 2500 platform with 50 bp paired end reads. Following quality assessment via FastQC, adapter and quality trimming was executed with Trim Galore. Reads were subsequently aligned to the *hg19* genome assembly via Bowtie2 with ~97% concordant alignment rate in all samples. After filtering for unmapped, low quality, and multi-mapped reads, mapped reads were summarized to gene features by HTSeq. Sequencing depth ranged from 30-42 million postfiltered reads. We used DESeq2 to evaluate significant cases of differential expression between a given pairwise comparison. Before adjusting p-values for multiple testing, DESeq2 implements independent filtering using mean expressions of each gene as a filter. Adjusting via the Benjamini & Hochberg method, differentially expressed genes with FDR < 0.05 were called significant. Prior to clustering, we used DESeq2’s implementation of regularized logarithm transformation (rlog) to stabilize the variance of genes across samples. Mean expression values were used as input to hierarchical clustering of the differentially expressed genes between soft v. stiff IPN. Gene ontology and TF association analysis using ChIP-seq data from ENCODE was implemented via EnrichR (9). Adjusted p-values, which take into account differing sizes of data sets, are reported.

#### Statistical Analysis

Multiple comparisons were conducted with one-way ANOVA followed by Tukey post-hoc comparison and pairwise comparisons performed using Student’s t-tests. Bars represent mean ± SEM and symbols represent each experiment (for EdU assays), cell cluster (for invasion assays), patient (for 3SEQ expression), and nucleus (for nuclear morphology assays). For graphs of RNA-seq normalized counts from 3D culture, bars represent mean ± SD and symbols represent each RNA-seq experiment using 2 independent trials. Values with p < 0.05 were considered statistically significant, indicated by **.

#### Data Availability

RNA-seq data are stored in GEO with the accession code GSE102506. 3SEQ breast cancer progression data are available as described in the original manuscript (10). Additional normal mammary and breast cancer datasets generated by the Genotype-Tissue Expression (GTEx) project and The Cancer Genome Atlas (TCGA) project, respectively, are available on the Human Protein Atlas (11, 12).

**Fig. S1.**
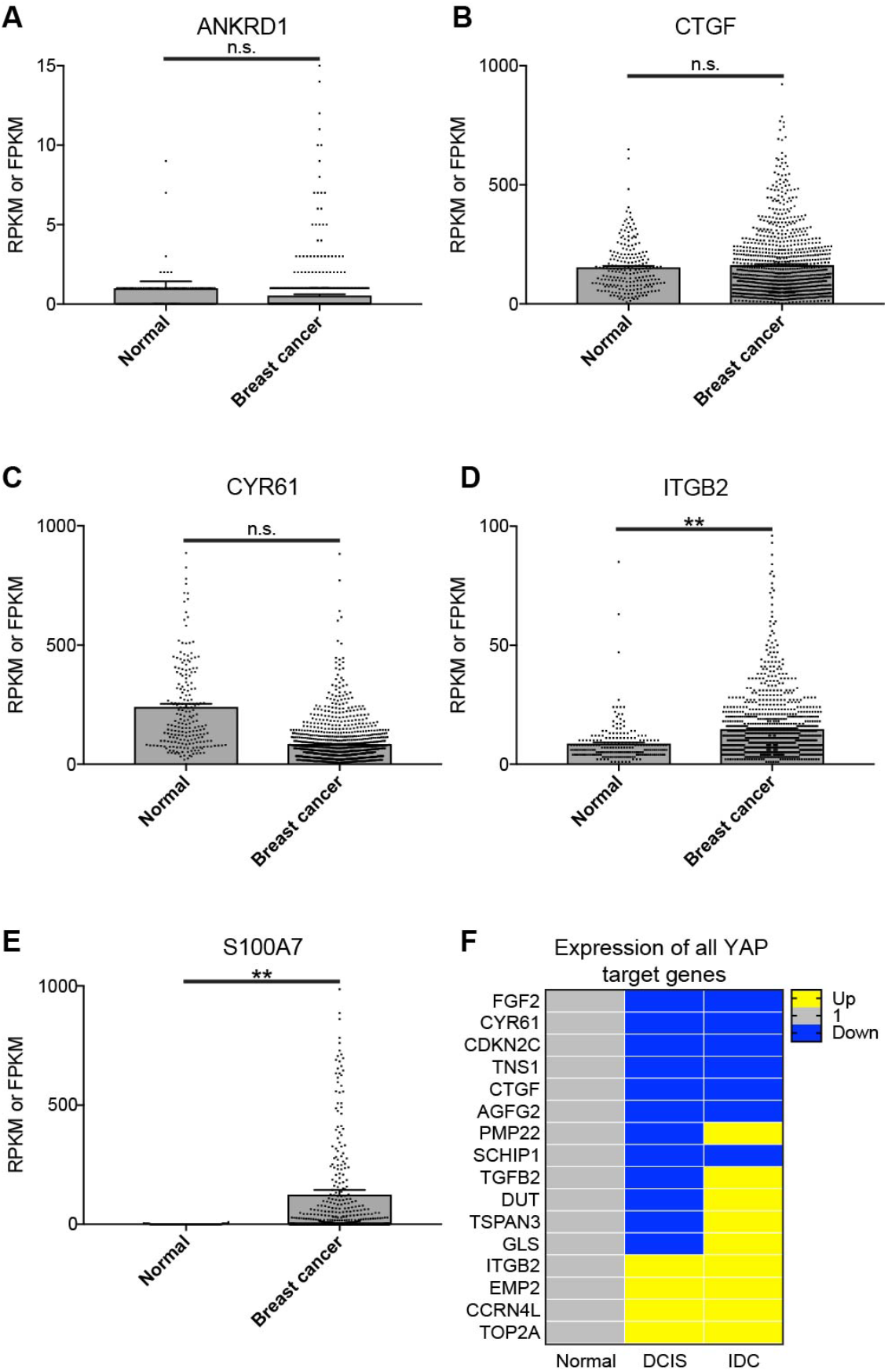
Expression of YAP target genes in breast cancer patient datasets. RNA-seq gene expression counts of canonical YAP target genes **(A)** ANKRD1, **(B)** CTGF, **(C)** CYR61, **(D)** ITGB2, and control **(E)** S100A7, in normal mammary and breast cancer tissues obtained from Human Protein Atlas {Uhlen:2015ip, Uhlen:2010eu}. Normal and breast cancer results are reported as RPKM (GTEx dataset) and FPKM (TCGA dataset), respectively. (n.s., p > 0. 05; **, p < 0.05, one-way ANOVA followed by Tukey post-hoc comparison tests, n = 214-1075 patient samples, error bars represent SEM, symbols represent each patient sample, some symbols beyond graph axes). **(F)** Diagram showing increased or decreased expression of YAP target genes during breast cancer progression from Brunner *et al.* study {Brunner:2014fk}.

**Fig. S2.**
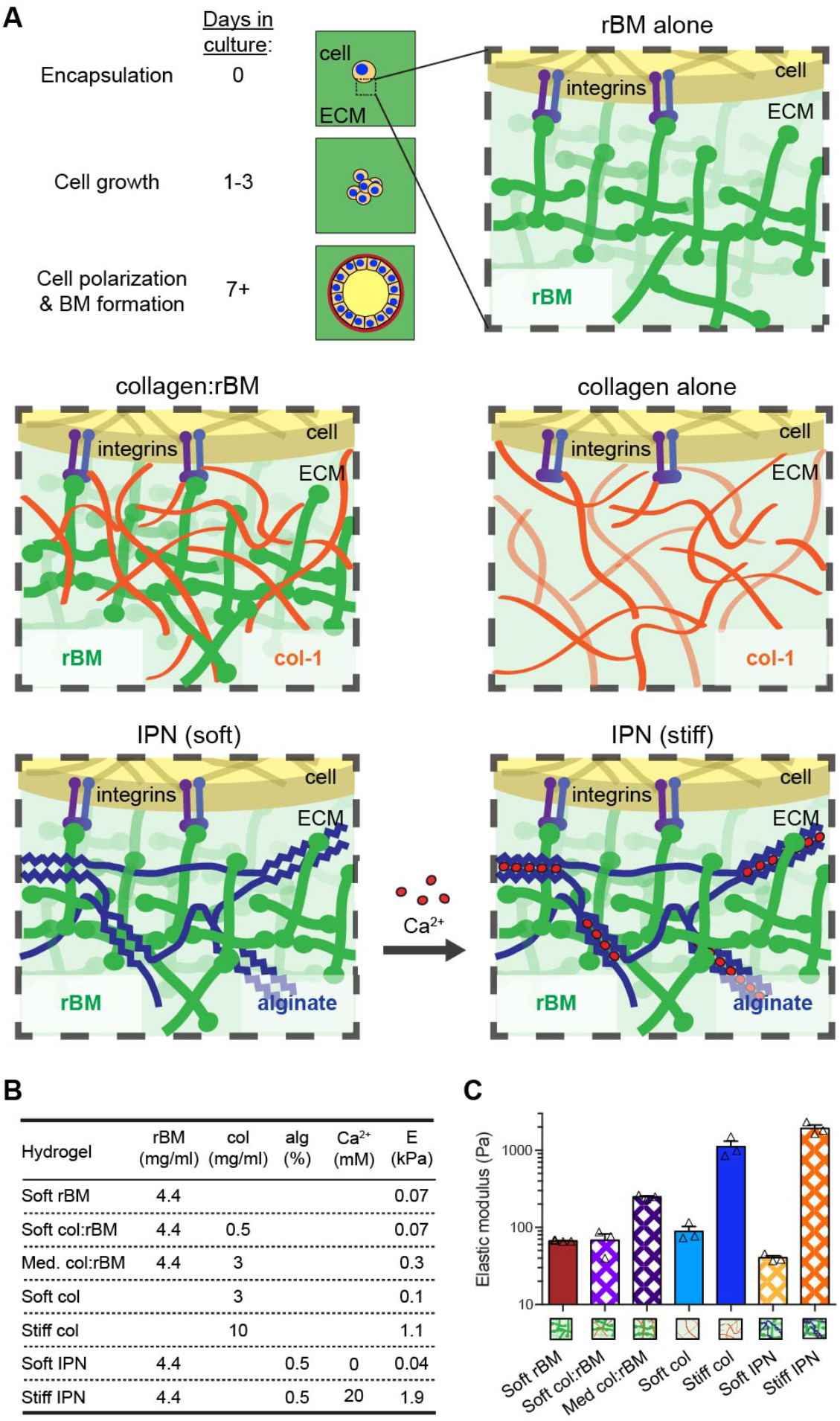
Generation of hydrogels with range of elastic moduli and col-1 concentration. **(A)** Schematic of hydrogel encapsulated MCF10A forming an acinar structure and hydrogels generated in this study. rBM: reconstituted Basement Membrane; col-1: type 1 collagen. **(B)** Composition and stiffness of hydrogels used in this study. alg: alginate; col: col-1; E: elastic modulus. **(C)** Elastic modulus measurements of hydrogels at 1 Hz. Bars represent mean of three gels ± SEM, symbols represent E of each gel.

**Fig. S3.**
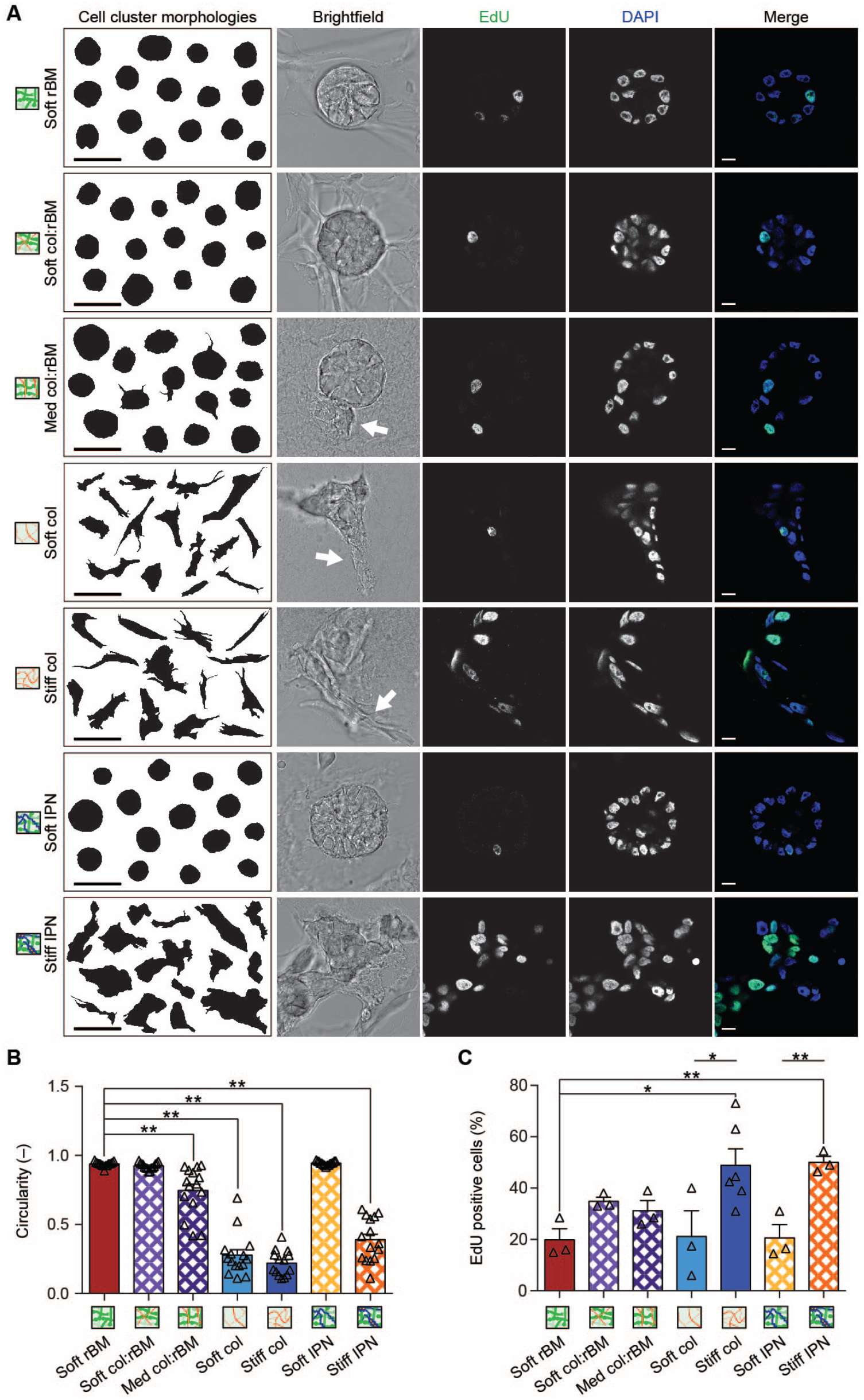
Enhanced ECM stiffness promotes invasion and proliferation in 3D culture in the presence or absence of col-1. **(A)** MCF10A cells encapsulated for seven days, with 24 h EdU treatment, were fixed and stained for EdU (green). DNA was stained using DAPI (blue), scale bars: 10 μm. Arrows denote invasive structures. Morphological analysis of 14-15 cell clusters from each hydrogel are displayed in first column, scale bars: 100 μm. **(B)** Invasiveness of cells in each hydrogel as measured by cell cluster circularity (**, p < 0.001, one-way ANOVA followed by Tukey post-hoc comparison tests, n = 14 cell clusters for stiff col, n = 15 cell clusters for all others, from 1 experiment, error bars represent SEM, symbols represent each cell cluster). **(C)** Graph of percent EdU positive cells in each hydrogel. (**, p < 0.05; *, p < 0.1, one-way ANOVA followed by Tukey post-hoc comparison tests, n = 200 cells per experiment, from 3 experiments for all except stiff col which had 6 experiments, error bars represent SEM, symbols represent mean of each experiment).

**Fig. S4.**
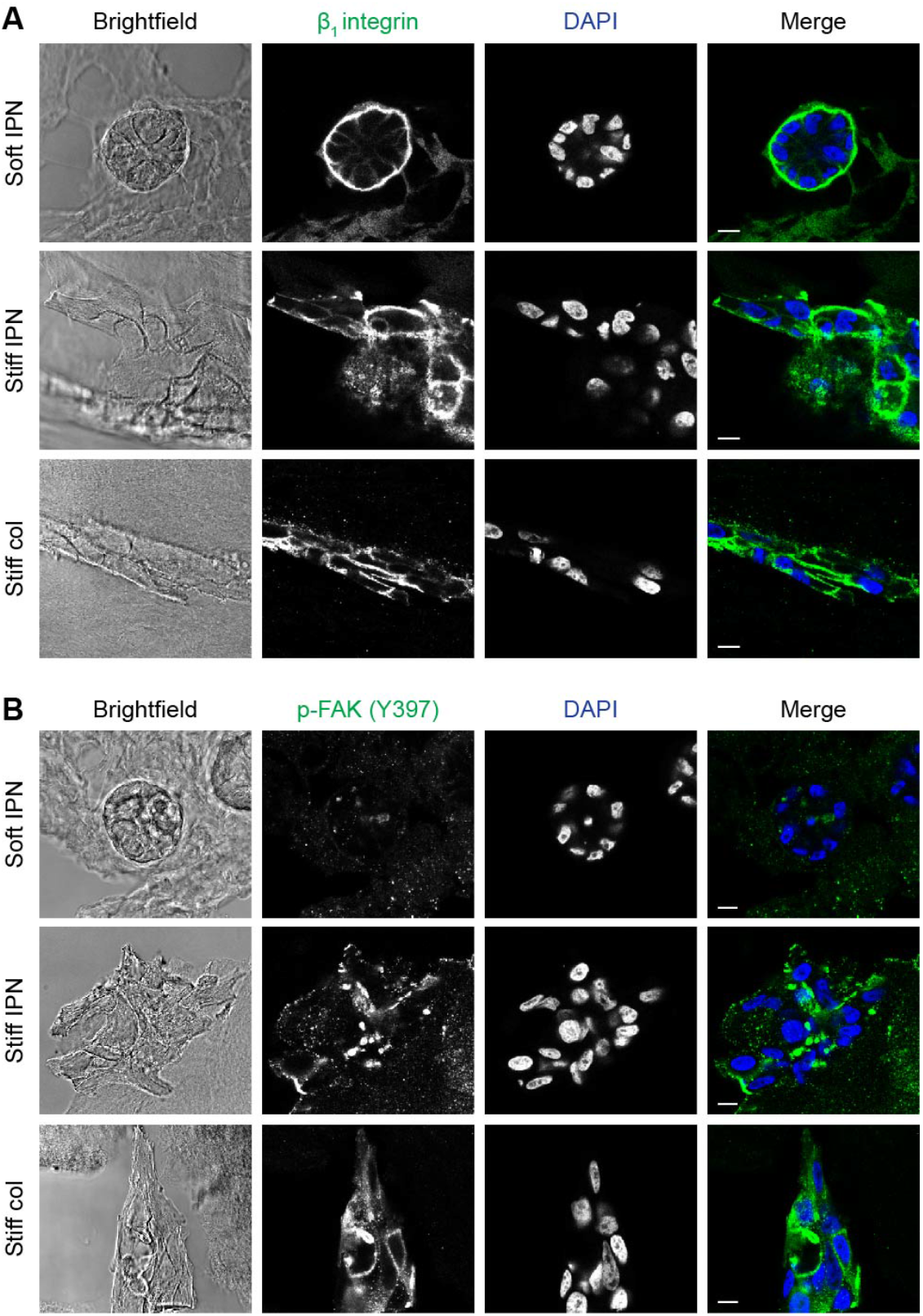
Assessment of general mechanotransduction pathways. Confocal micrographs of MCF10A cells encapsulated for seven days in indicated hydrogels and stained for **(a)** β_1_ integrin (green) or **(b)** phospho-FAK (Y397) (green). DNA is stained using DAPI (blue). Scale bars are 10 μm.

**Fig. S5.**
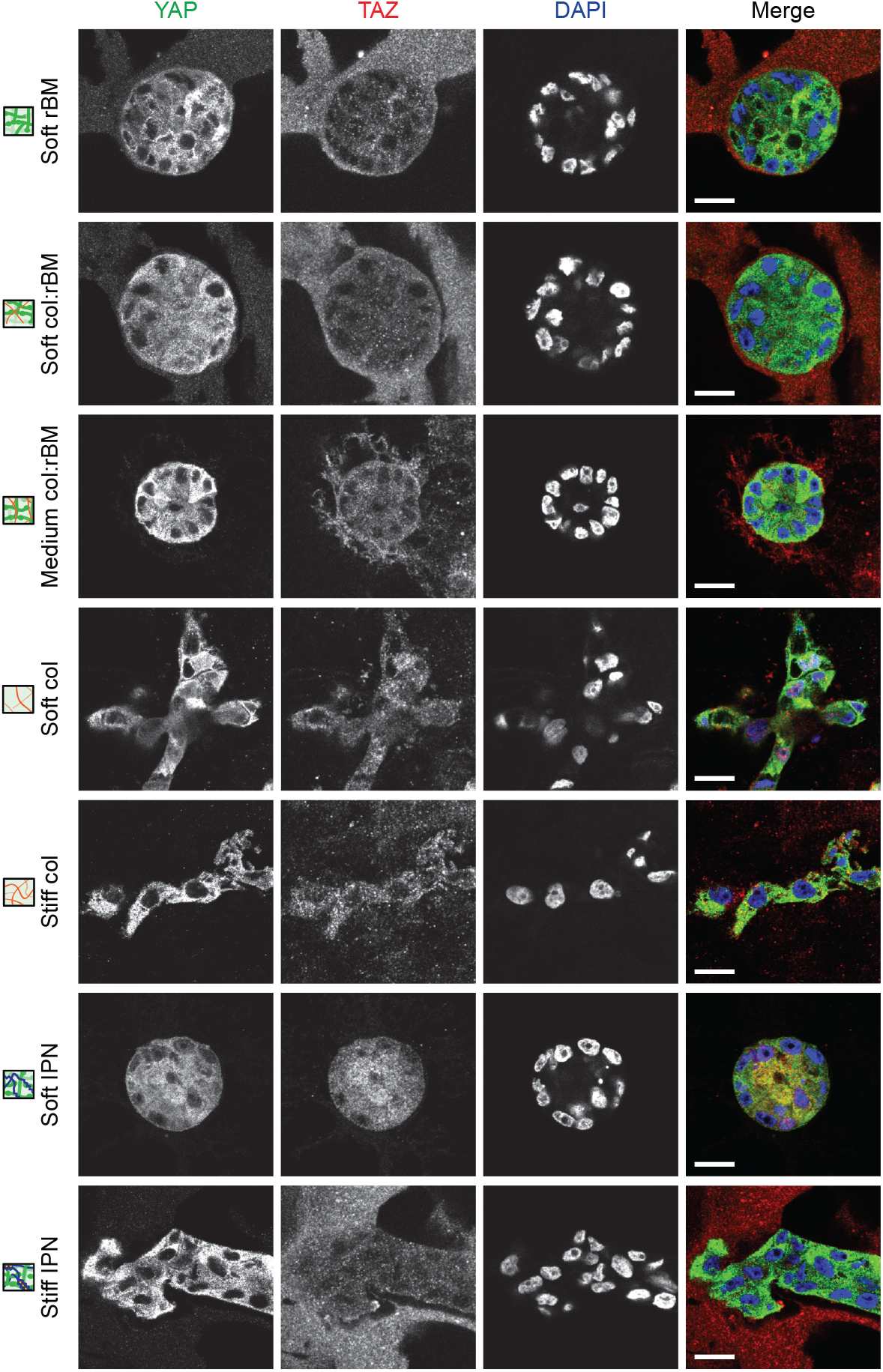
TAZ localization. Confocal micrographs of MCF10A cells encapsulated for seven days in indicated hydrogels corresponding to Fig. 2 stained for YAP (green), TAZ (red), and DNA using DAPI (blue). Scale bars: 10 μm. We note there is also ECM staining by the TAZ antibody.

**Fig. S6.**
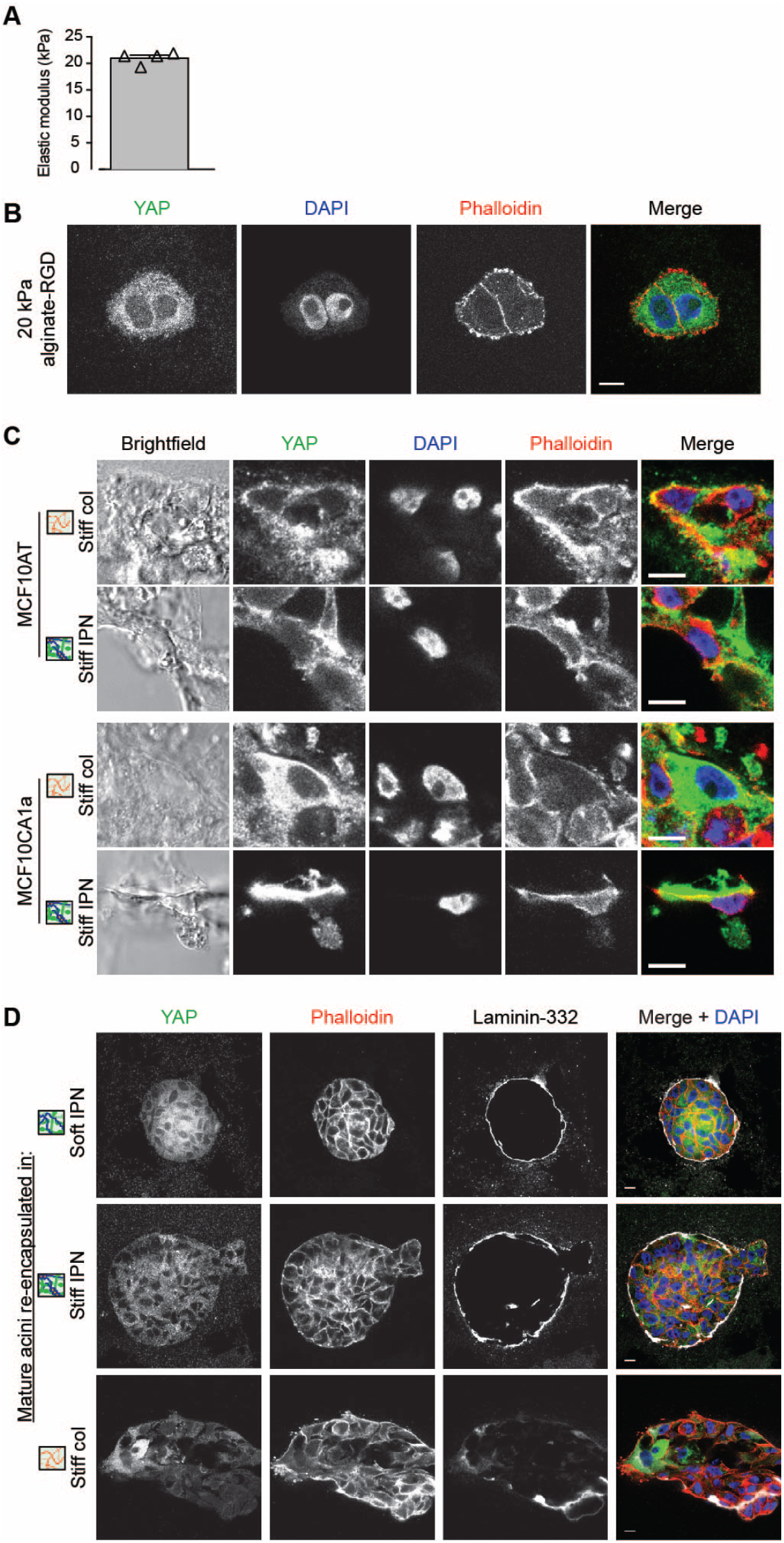
YAP staining with 20 kPa stiffness, MCF10AT and MCF10CA1a cells, and transplantations. **(A)** Unconfined compression of hydrogels compressed to 15% at a rate of 1 mm/min to obtain the stiffness of alginate hydrogels. Bars represent mean of four gels ± SEM, symbols represent E of each gel. **(B)** MCF10A cells encapsulated for three days in 20 kPa alginate-RGD. **(C)** MCF10AT and MCF10CA1a cells encapsulated for seven days. YAP (green). F-actin was stained with phalloidin (red) and DNA with DAPI (blue). Scale bars: 10 μm. **(D)** Mature MCF10A acini harvested from seven day rBM overlay cultures (Debnath et al., 2003) and then encapsulated in stiff IPNs for three days. Cells were stained for YAP (green), F-actin using phalloidin (red), laminin-332 (white), and DNA using DAPI (blue). Scale bar: 10 μm.

**Fig. S7.**
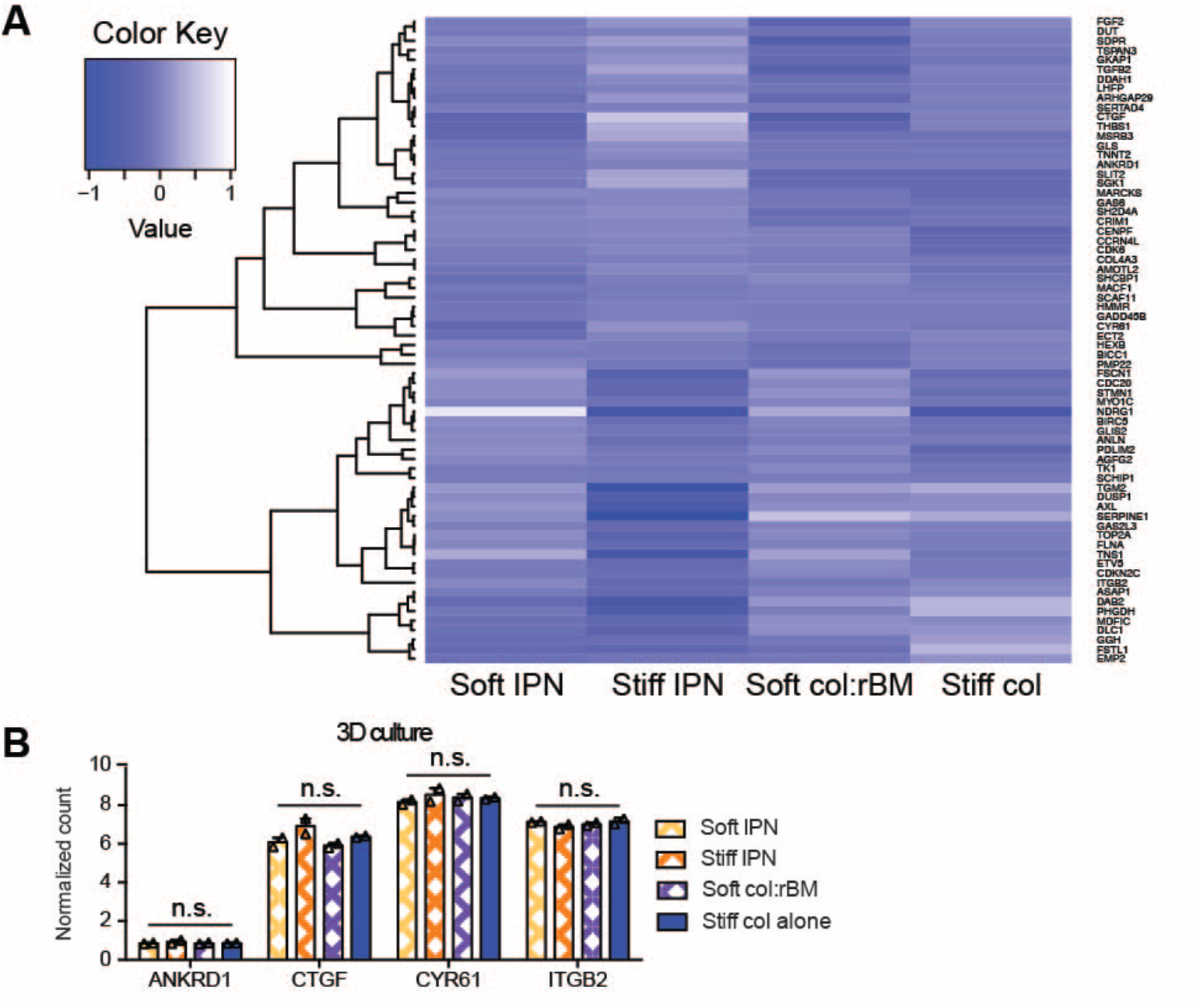
Enhanced col-1 stiffness does not promote YAP target gene expression. **(A)** Expression of YAP target genes (as identified by Dupont et al., 2011) in MCF10A cells encapsulated in indicated hydrogels for seven days and analyzed by RNA-seq. Values are normalized by mean of each row. **(B)** Graph of RNA-seq normalized counts of canonical YAP target genes in each hydrogel. (n.s., p > 0.05; one-way ANOVA followed by Tukey post-hoc comparison tests, n = 2 independent experiments, error bars represent S.D., symbols represent each experiment).

**Fig. S8.**
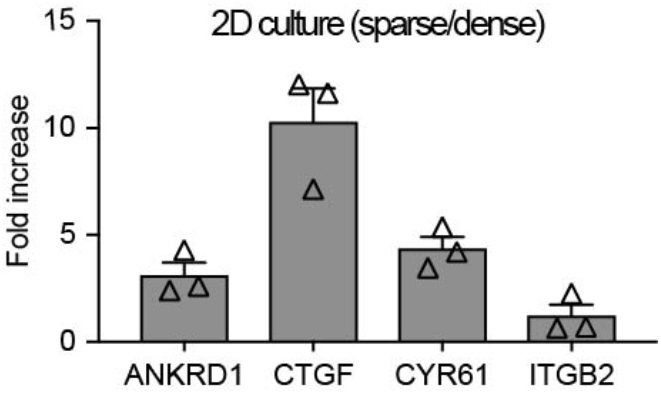
YAP target gene expression in 2D culture. Graph of expression fold change of canonical YAP target genes with YAP activating (2D sparse plating) over YAP inactivating (2D dense plating) conditions in MCF10A cells. (n = 3 samples from 1 experiment, error bars represent SEM, symbols represent each replicate).

**Fig. S9.**
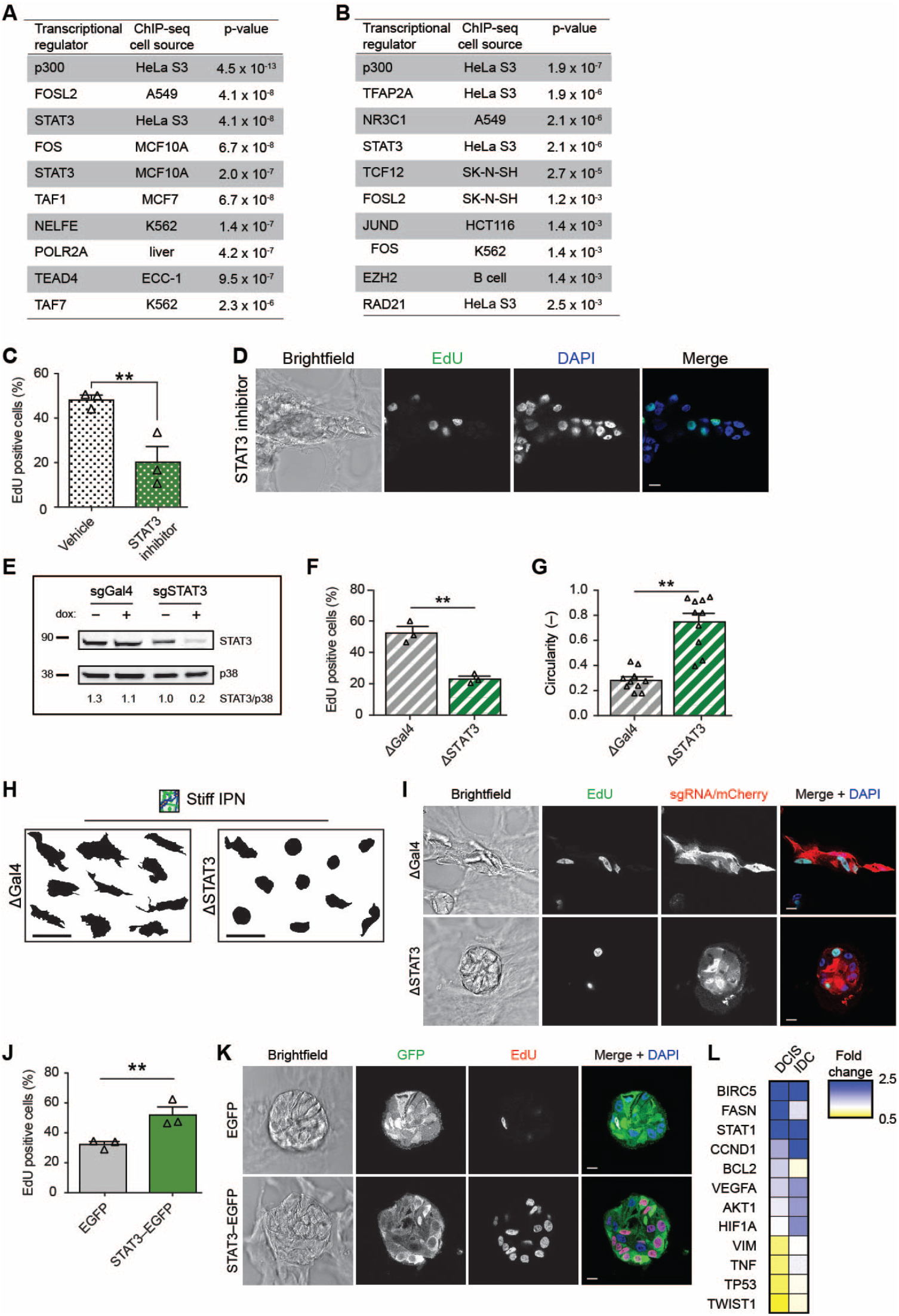
STAT3-dependent mechanotransduction in 3D culture. Table of transcriptional regulators most significantly associated with DE genes induced by increased **(A)** IPN stiffness (stiff IPN/soft IPN) or **(B)** col-1 stiffness (stiff col/soft col:rBM), as defined by ChIP-seq data provided by ENCODE. **(C)** Graph of percent EdU positive MCF10A cells encapsulated for seven days in stiff IPNs and treated for three days with 500 μM STAT3 peptide inhibitor PY*LKTK or vehicle (DMSO). (**, p < 0.05, one-way ANOVA followed by Tukey post-hoc comparison tests, n = 200 cells per experiment, from 3 experiments, error bars represent SEM, symbols represent mean of each experiment). **(D)** Confocal images of encapsulated MCF10A cells from (C) stained for EdU (green) and DNA using DAPI (blue). Scale bars: 10 μm. **(E)** Western blot analysis of MCF10A::Cas9/sgGAL4 or MCF10A::Cas9/sgSTAT3 cells with or without 2 μg/ml dox for 72 h and probed for STAT3. p38 was used as a loading control. Fluorescence intensity quantification of bands, normalized by p38 control, are indicated below each lane. **(F)** Graph of percent EdU positive cells in each hydrogel. (**, p < 0.01, one-way ANOVA followed by Tukey post-hoc comparison tests, n = 200 cells per experiment, from 3 experiments, error bars represent SEM, symbols represent mean of each experiment). **(G)** Invasiveness of cells in each hydrogel as measured by cell cluster circularity, (**, p < 0.01, oneway ANOVA followed by Tukey post-hoc comparison tests, n = 10 cell clusters, from 1 experiment, error bars represent SEM, symbols represent each cell cluster). **(H)** Cell cluster morphologies of ΔGAL4 (control) or ΔSTAT3 MCF10A cells encapsulated for seven days with 2 μg/ml dox in stiff IPNs. Scale bars: 100 μm. **(I)** Representative cluster from (H) stained for EdU (green), sgRNA/mCherry (red), and DNA using DAPI (blue). Scale bars: 10 μm. **(J)** Graph of percent EdU positive MCF10A cells transduced with STAT3-EGFP or EGFP control and encapsulated for seven days in stiff IPNs from ( **, p < 0.01, one-way ANOVA followed by Tukey post-hoc comparison tests, n = 200 cells per experiment, from 3 experiments, error bars represent SEM, symbols represent mean of each experiment). **(K)** Representative image from (J) stained for GFP (green), EdU (red), and DNA using DAPI (blue). Scale bars: 10 μm. **(L)** Heat map showing expression of selected STAT3 target genes that are highly relevant to cancer in DCIS or IDC patient samples. Values are displayed as fold change compared to normal.

**Fig. S10.**
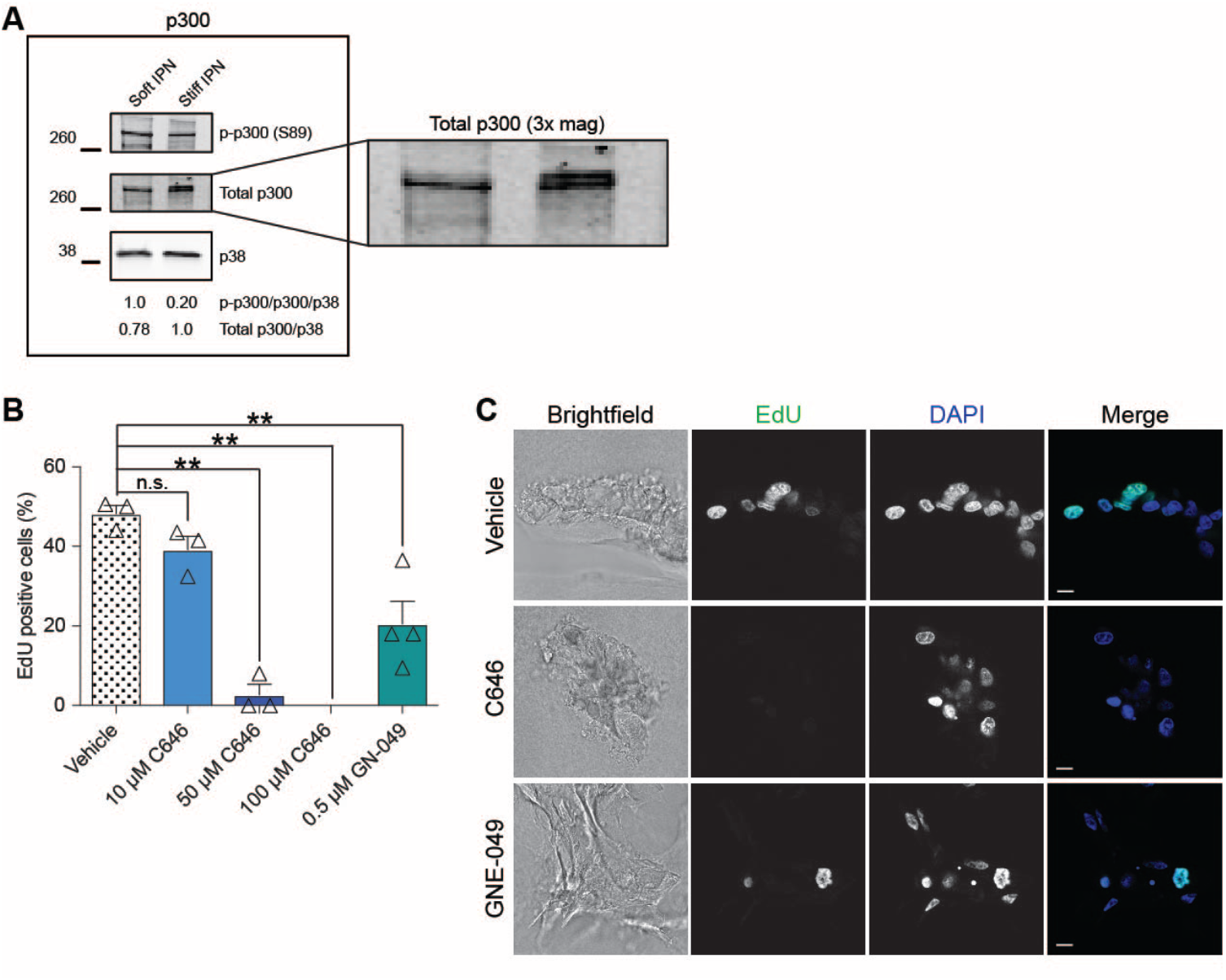
p300 stiffness-induced gene regulation. **(A)** Western blot analysis of cells harvested from soft and stiff IPNs and probed for p-p300 (S89) and total p300 (with 3x magnification) p38 was used as a loading control. Fluorescence intensity quantification of bands, normalized by p38 control, are indicated below each lane. **(B)** Graph of percent EdU positive MCF10A cells encapsulated for seven days in stiff IPNs and treated for three days with p300 inhibitors C646 or GNE-049 or vehicle (DMSO). (n.s., p > 0.05; **, p < 0.05, one-way ANOVA followed by Tukey post-hoc comparison tests, n = 200 cells per experiment, from 3 experiments, error bars represent SEM, symbols represent mean of each experiment). **(C)** Confocal images of 100 μM C646 or 0.5 μM GNE-049 treated cells from (e) stained for EdU (green) and DNA using DAPI (blue). Scale bars: 10 μm.

**Fig. S11.**
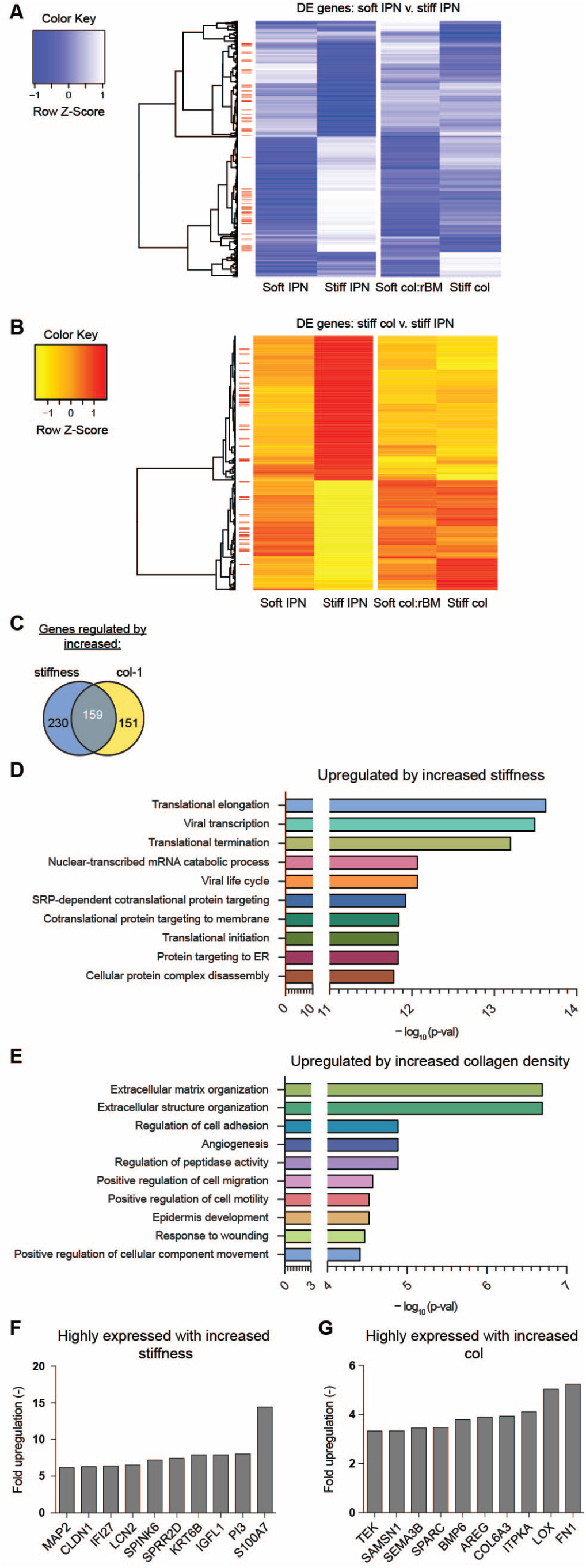
RNA-seq analysis reveals alteration in gene expression of MCF10A cells due to increased stiffness or enhanced collagen density in 3D culture. **(A)** Heat map of 389 genes differentially expressed (DE) at FDR < 0.05 when comparing MCF10A cells harvested from soft v. stiff IPNs. Red bars denote genes involved in proliferation, cell cycle, mitosis, or adhesion. Values are displayed as row Z-scores. **(B)** Heat map of 310 genes DE at FDR < 0.05 when comparing stiff IPNs v. stiff col-1 hydrogels. **(C)** Venn diagram showing number of genes regulated by both enhanced stiffness and enhanced col-1 density. **(D)** Bar graph of gene ontology (GO) terms significantly represented in differentially expressed genes during increased stiffness (stiff IPN/soft IPN) or **(E)** increased col-1 (stiff col/stiff IPN). **(F)** Bar graph of most highly upregulated genes during increased stiffness or **(G)** increased col-1.

**Fig. S12.**
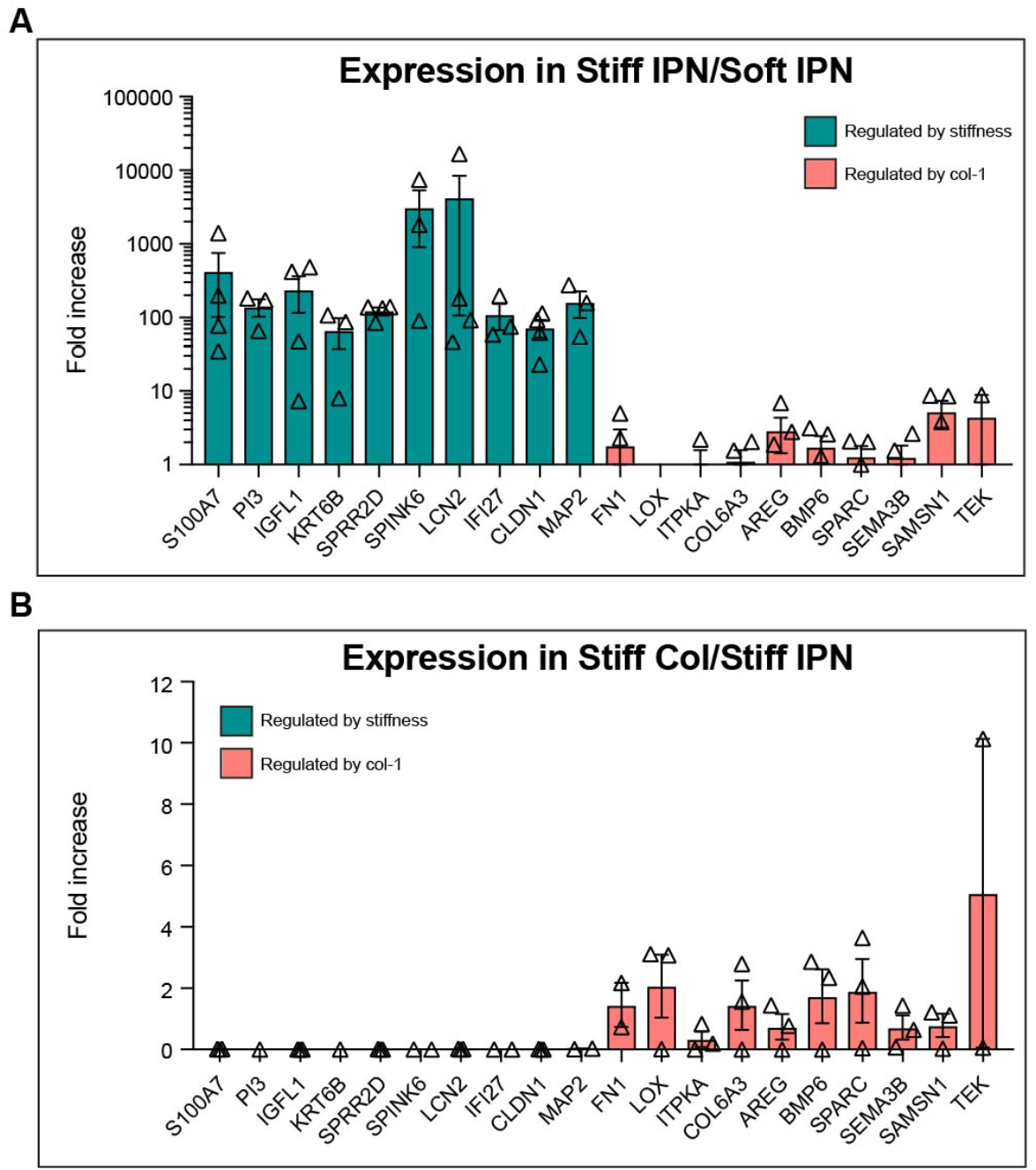
qPCR validation of 3D culture RNA-seq. **(A)** Graph showing qPCR results of RNA-seq identified stiffness- and col-regulated genes in 3D cultured stiff IPNs (compared to soft IPNs). **(B)** Graph showing qPCR results of RNA-seq identified stiffness- and col-regulated genes in 3D cultured stiff col-1 hydrogels (compared to stiff IPNs). (n = 3 experiments, error bars represent SEM, symbols represent each experiment).

**Fig. S13.**
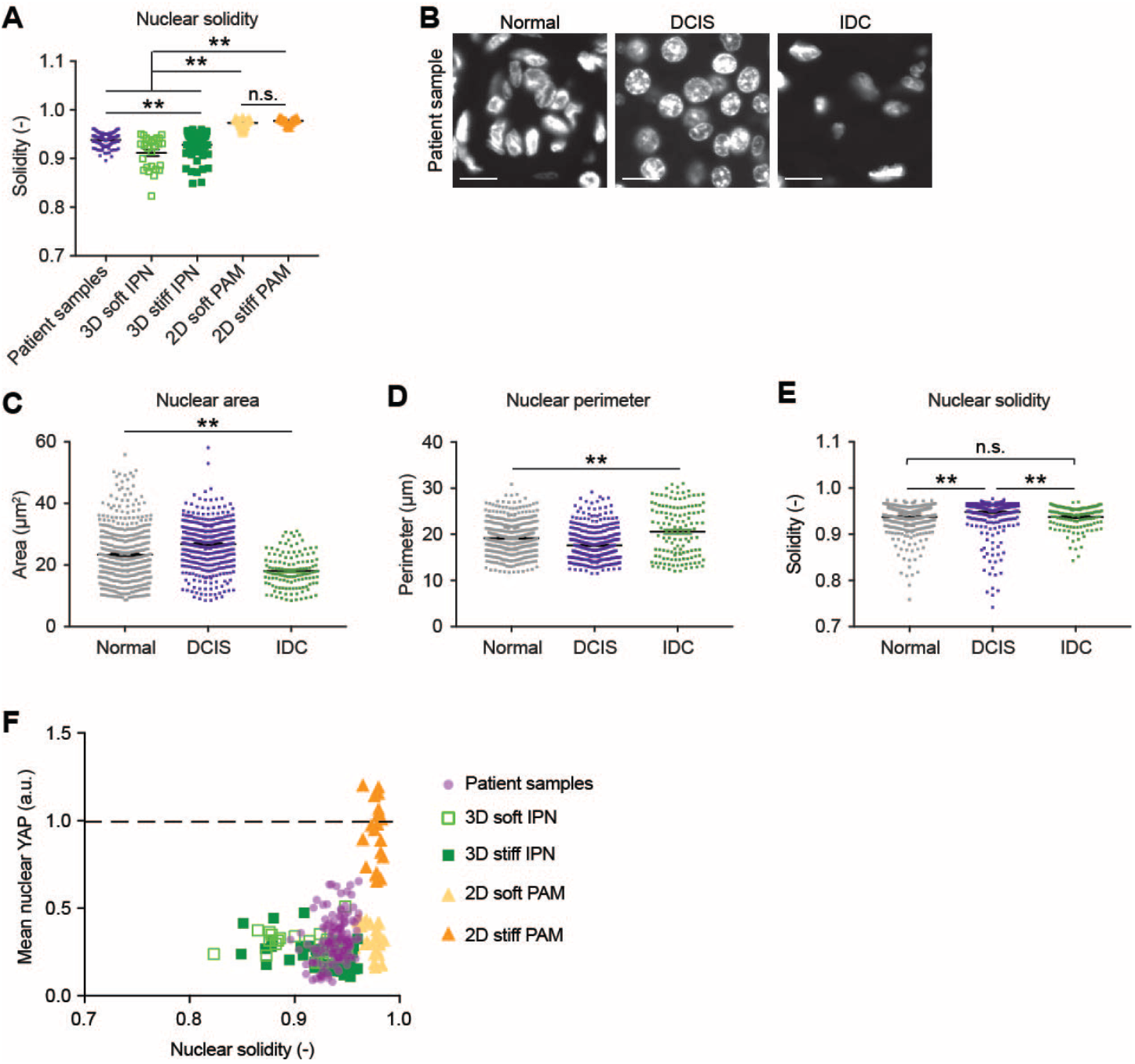
Nuclear solidity and morphology changes during *in vivo* cancer progression. **(A)** Solidity of nuclei (value of 1 represents a perfect circle). Patient samples from five DCIS patients. **(B)** Images of nuclear morphologies during breast cancer progression. Bars: 10 μm. **(C)** Areas, **(B)** perimeters, and **(C)** solidity of nuclei. Patient samples from five normal, 5 DCIS, and 5 IDC patients. **(F)** YAP intensity with nuclear solidity.

**Table S1.**
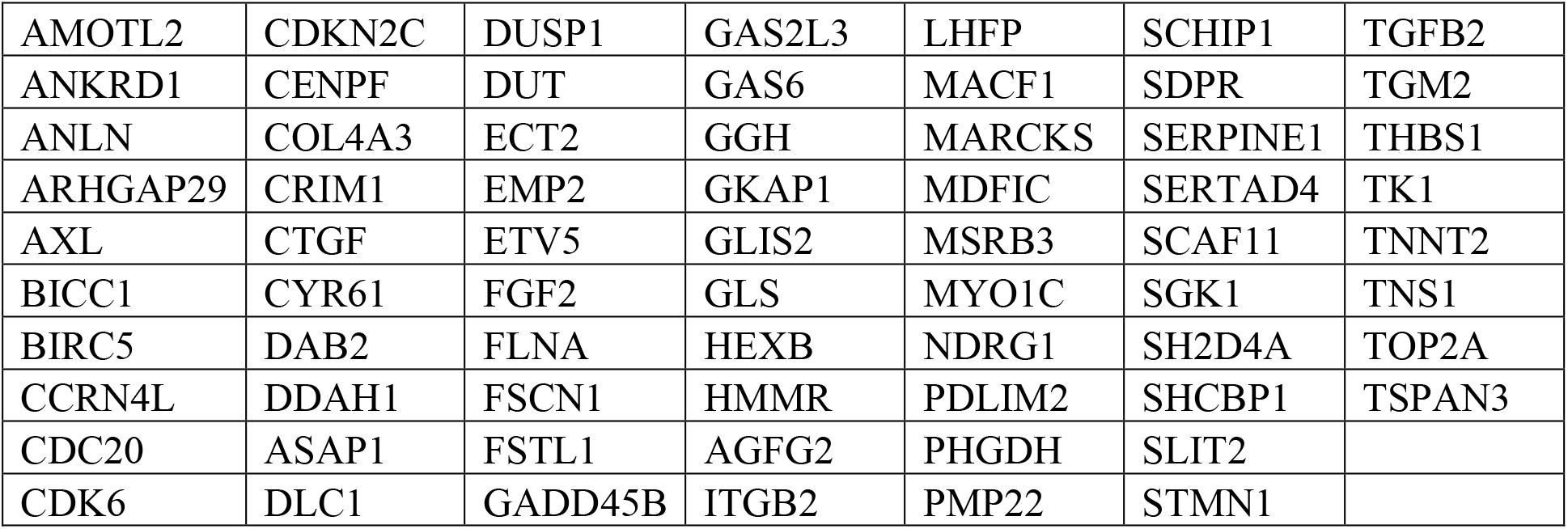
YAP target genes used in Fig. 2F.

**Table S2.**
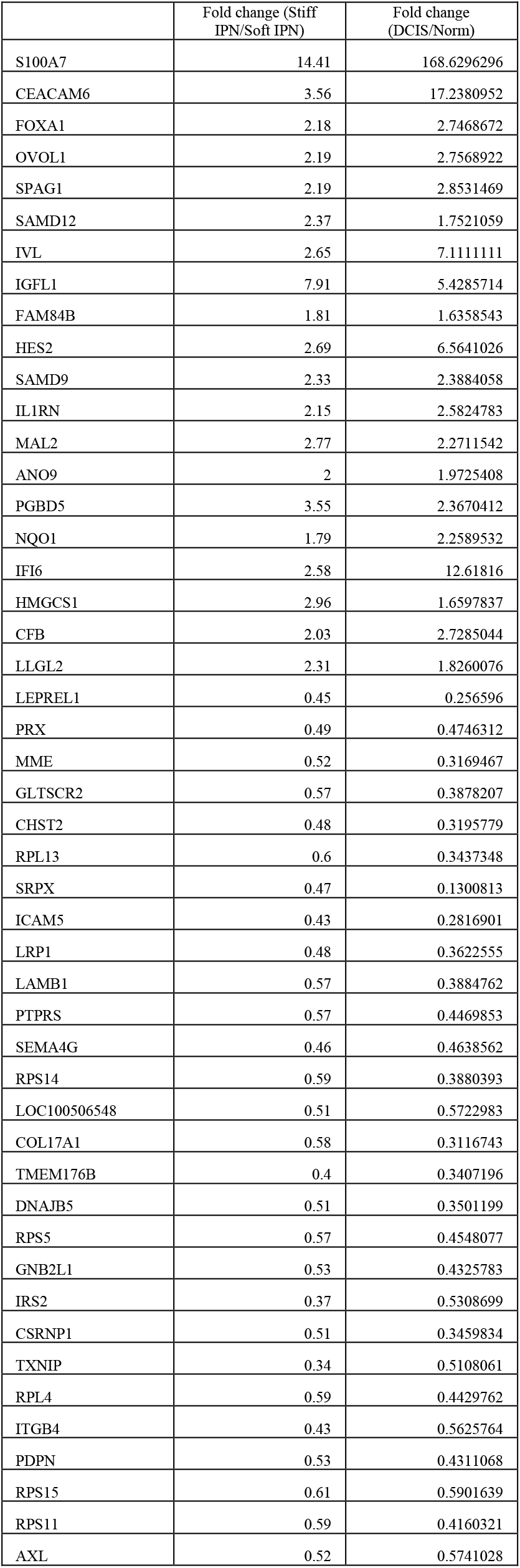
Genes with similar regulation in stiff IPNs as DCIS samples in Fig. 2L.

## References

1. N. F. Boyd, G. A. Lockwood, J. W. Byng, D. L. Tritchler, M. J. Yaffe, Mammographic densities and breast cancer risk. Cancer Epidemiol. Biomarkers Prev. 7, 1133–1144 (1998).

2. L. J. Martin, N. F. Boyd, Mammographic density. Potential mechanisms of breast cancer risk associated with mammographic density: hypotheses based on epidemiological evidence. Breast Cancer Res. 10, 201 (2008).

3. J. C. McConnell et al., Increased peri-ductal collagen micro-organization may contribute to raised mammographic density. Breast Cancer Res. 18, 5 (2016).

4. K. R. Levental etal., Matrix crosslinking forces tumor progression by enhancing integrin signaling. Cell. 139, 891–906 (2009).

5. I. Acerbi et al., Human breast cancer invasion and aggression correlates with ECM stiffening and immune cell infiltration. Integr Biol (Camb). 7, 1120–1134 (2015).

6. R. Sinkus et al., MR elastography of breast lesions: understanding the solid/liquid duality can improve the specificity of contrast-enhanced MR mammography. Magn Reson Med. 58, 1135–1144 (2007).

7. M. J. Paszek et al., Tensional homeostasis and the malignant phenotype. Cancer Cell. 8, 241–254 (2005).

8. P. P. Provenzano et al., Collagen density promotes mammary tumor initiation and progression. BMC Med. 6, 11 (2008).

9. S. C. Wei etal., Matrix stiffness drives epithelial-mesenchymal transition and tumour metastasis through a TWIST1-G3BP2 mechanotransduction pathway. Nat. Cell Biol. 17, 678–688 (2015).

10. O. Chaudhuri etal., Extracellular matrix stiffness and composition jointly regulate the induction of malignant phenotypes in mammary epithelium. Nat Mater. 13, 970–978 (2014).

11. P. D. Yurchenco, Basement membranes: cell scaffoldings and signaling platforms. Cold Spring Harb Perspect Biol. 3 (2011), doi:10.1101/cshperspect.a004911.

12. A. Totaro, T. Panciera, S. Piccolo, YAP/TAZ upstream signals and downstream responses. Nat. Cell Biol. 20, 888–899 (2018).

13. S. Dupont etal., Role of YAP/TAZ in mechanotransduction. Nature. 474, 179–183 (2011).

14. R. Oria etal., Force loading explains spatial sensing of ligands by cells. Nature. 552, 219–224 (2017).

15. A. Elosegui-Artola etal., Force Triggers YAP Nuclear Entry by Regulating Transport across Nuclear Pores. Cell (2017), doi:10.1016/j.cell.2017.10.008.

16. J.-Y. Shiu, L. Aires, Z. Lin, V. Vogel, Nanopillar force measurements reveal actin-cap-mediated YAP mechanotransduction. Nat. Cell Biol. 20, 262–271 (2018).

17. C. Fischbach etal., Engineering tumors with 3D scaffolds. Nat Meth. 4, 855–860 (2007).

18. E. Cukierman, R. Pankov, D. R. Stevens, K. M. Yamada, Taking cell-matrix adhesions to the third dimension. Science. 294, 1708–1712 (2001).

19. O. W. Petersen, L. Rønnov-Jessen, A. R. Howlett, M. J. Bissell, Interaction with basement membrane serves to rapidly distinguish growth and differentiation pattern of normal and malignant human breast epithelial cells. Proc. Natl. Acad. Sci. U.S.A. 89, 9064–9068 (1992).

20. J. Y. Lee, O. Chaudhuri, Regulation of breast cancer progression by extracellular matrix mechanics: insights from 3D culture models. ACS Biomaterials Science & Engineering (2017), doi:10.1021/acsbiomaterials.7b00071.

21. D. Elster etal., TRPS1 shapes YAP/TEAD-dependent transcription in breast cancer cells. Nat Commun. 9, 3115 (2018).

22. B. von Eyss, etal., A MYC-Driven Change in Mitochondrial Dynamics Limits YAP/TAZ Function in Mammary Epithelial Cells and Breast Cancer. Cancer Cell. 28, 743–757 (2015).

23. S. A. S. Real etal., Aberrant Promoter Methylation of YAP Gene and its Subsequent Downregulation in Indian Breast Cancer Patients. BMC Cancer. 18, 711 (2018).

24. H. Gao etal., Multi-organ Site Metastatic Reactivation Mediated by Non-canonical Discoidin Domain Receptor 1 Signaling. Cell. 166, 47–62 (2016).

25. J. Kiefer, A. Alexander, M. C. Farach-Carson, Type I collagen-mediated changes in gene expression and function of prostate cancer cells. Cancer Treat. Res. 118, 101–124 (2004).

26. S. J. Santner et al., Malignant MCF10CA1 cell lines derived from premalignant human breast epithelial MCF10AT cells. Breast Cancer Res. Treat. 65, 101–110 (2001).

27. M. Aragona etal., A mechanical checkpoint controls multicellular growth through YAP/TAZ regulation by actin-processing factors. Cell. 154, 1047–1059 (2013).

28. A. L. Brunner et al., A shared transcriptional program in early breast neoplasias despite genetic and clinical distinctions. Genome Biol. 15, R71 (2014).

29. E. D. Emberley, L. C. Murphy, P. H. Watson, S100A7 and the progression of breast cancer. Breast Cancer Res. 6, 153–159 (2004).

30. M. W. Nasser et al., RAGE mediates S100A7-induced breast cancer growth and metastasis by modulating the tumor microenvironment. Cancer Res. 75, 974–985 (2015).

31. S. Mandal, L. Curtis, M. Pind, L. C. Murphy, P. H. Watson, S100A7 (psoriasin) influences immune response genes in human breast cancer. Exp. Cell Res. 313, 3016–3025 (2007).

32. S. Lee, S. Kumar, Actomyosin stress fiber mechanosensing in 2D and 3D. F1000Res. 5, 2261 (2016).

33. Y. A. Miroshnikova etal., Adhesion forces and cortical tension couple cell proliferation and differentiation to drive epidermal stratification. Nat. Cell Biol. 20, 69–80 (2018).

## References

1. J. Debnath, S. K. Muthuswamy, J. S. Brugge, Morphogenesis and oncogenesis of MCF10A mammary epithelial acini grown in three-dimensional basement membrane cultures. Methods. 30, 256–268 (2003).

2. O. Chaudhuri et al., Extracellular matrix stiffness and composition jointly regulate the induction of malignant phenotypes in mammary epithelium. Nat Mater. 13, 970–978 (2014).

3. O. Chaudhuri et al., Substrate stress relaxation regulates cell spreading. Nat Commun. 6, 6364 (2015).

4. J. A. Rowley, G. Madlambayan, D. J. Mooney, Alginate hydrogels as synthetic extracellular matrix materials. Biomaterials. 20, 45–53 (1999).

5. R. S. Fischer, K. A. Myers, M. L. Gardel, C. M. Waterman, Stiffness-controlled three-dimensional extracellular matrices for high-resolution imaging of cell behavior. Nat Protoc. 7, 2056–2066 (2012).

6. J. R. Tse, A. J. Engler, Preparation of hydrogel substrates with tunable mechanical properties. Curr Protoc Cell Biol. Chapter 10, Unit 10.16 (2010).

7. K. R. Levental et al., Matrix crosslinking forces tumor progression by enhancing integrin signaling. Cell. 139, 891–906 (2009).

8. M. J. Paszek et al., Tensional homeostasis and the malignant phenotype. Cancer Cell. 8, 241–254 (2005).

9. M. V. Kuleshov et al., Enrichr: a comprehensive gene set enrichment analysis web server 2016 update. Nucleic Acids Res. 44, W90–7 (2016).

10. A. L. Brunner et al., A shared transcriptional program in early breast neoplasias despite genetic and clinical distinctions. Genome Biol. 15, R71 (2014).

11. M. Uhlén et al., Towards a knowledge-based Human Protein Atlas. Nat. Biotechnol. 28, 1248–1250 (2010).

12. M. Uhlén et al., Proteomics. Tissue-based map of the human proteome. Science. 347, 1260419 (2015).

